# On the synergy between myelin proteins P0, MBP, and P2 in peripheral nerve major dense line formation

**DOI:** 10.1101/2024.07.15.603506

**Authors:** Oda C. Krokengen, Arne Raasakka, Martin Berg Klenow, Antara Pal, Øystein Hetland, Anna Mularski, Salla Ruskamo, Jan Skov Pedersen, Adam Cohen Simonsen, Petri Kursula

## Abstract

Myelin is a proteolipid membrane multilayer held together by a set of proteins. The proper formation and function of the myelin sheath relies on the coordinated action of several key myelin proteins. Research exploring how proteins from the peripheral myelin cytoplasmic apposition – myelin basic protein (MBP), the cytoplasmic tail of myelin protein zero (P0ct), and peripheral myelin protein 2 (P2) – interact with each other and with myelin-like membranes was conducted using various techniques, such as small-angle X-ray diffraction (SAXD), differential scanning calorimetry (DSC), surface plasmon resonance (SPR), as well as electron and live epifluorescence microscopy. DSC revealed changes in lipid interactions depending on the protein combination, with MBP and P0ct binding more tightly to lipid membranes than P2, resulting in altered membrane fluidity and stability. These results were supported by SPR, which indicated that the myelin proteins may compete for membrane surface binding. Analysis of the Bragg peaks induced by the myelin proteins in lipidic environments showed both lamellar and non-lamellar phases in protein-lipid complexes. The results indicate both synergy and competition between the three main proteins residing in the PNS myelin major dense line. Furthermore, the observed direct effects of myelin proteins on lipid membrane properties may be relevant to their function in myelinating cells.

## Introduction

The correct structure and function of the myelin sheath depend on intricate relationships between the multiple layers of a lipid membrane and a unique group of proteins. Whereas a typical cell membrane has an approximate protein-to-lipid mass ratio of 1:1, myelin harbors a high proportion of lipids (70-80%) and a low amount of protein (15-30%) [1]. Formation of the substructure compact myelin (CM) and the periodical organization of the major dense line (MDL) and the intraperiod line (IPL) is vital for proper saltatory conduction of nerve impulses [2]. This closely packed structure is held together by amphipathic lipids and membrane-bound proteins that exert repulsive forces toward the surrounding liquid [2]. Even though the lipid content of CM is similar in both the central (CNS) and peripheral nervous system (PNS), the protein composition differs [1, 2].

Two major groups of proteins occupy ∼90% of the total PNS myelin protein content: glycoproteins and basic proteins. The three most abundant proteins in PNS CM are myelin protein zero (P0), myelin basic protein (MBP), and peripheral myelin protein 2 (P2) [3]; more than 50% of the total protein is represented by the 30-kDa glycoprotein P0 [4]. P0 is a transmembrane protein carrying an Ig-like domain at the IPL that is connected to a basic 69-residue C-terminal cytoplasmic tail (P0ct) through a single transmembrane α-helix [5]. This highly conserved protein is believed to function as an adhesion molecule by a head-to-head oligomerization of apposing Ig domains [6, 7] and electrostatic interactions between its tail and negatively charged lipids [8]. The intrinsically disordered P0ct is regulated by post-translational modifications, and a compromised cytoplasmic tail affects the function of the full-length protein as an adhesive molecule [9, 10]. Multiple mutations in P0 cause human neurological diseases, including Charcot-Marie-Tooth disease (CMT) [11–15], Dejerine-Sottas syndrome [16, 17], and congenital hypomyelination [18].

While MBP is the major protein in CNS myelin, occupying ∼30% of total myelin protein [19], it is less abundant in the PNS [4]. This disordered protein interacts with the phospholipid-rich face of the myelin membrane, resulting in MBP folding and partial membrane insertion within the MDL [20]. The 18.5-kDa highly basic main isoform of MBP is most abundant in healthy myelin, but various post-translational modifications, such as phosphorylation, deamidation, and methylation, result in a total of eight charge isoforms (C1-C8) with decreasing net positive charge affecting lipid binding [21, 22]. MBP is associated with multiple sclerosis and demyelination of the CNS [23], and although it plays a crucial role in the CNS, PNS compact myelin appears normal in MBP-deficient mice [24]. Its exact role in PNS myelin is unclear; besides being an adhesion molecule in CM, MBP interacts with other proteins [25], potentially guiding the number of Schmidt-Lanterman incisures and regulating the expression of connexin-32 (Cx32) and myelin-associated glycoprotein (MAG) [24].

In contrast to the intrinsically disordered P0ct and MBP, P2 is a small 15-kDa β-barrel with an α-helical lid, belonging to the family of fatty acid-binding proteins (FABP) [26, 27]. Two arginine residues, Arg106 and Arg126, inside the ligand binding pocket of the β-barrel, and positive patches on the P2 surface are essential for interacting with negatively charged lipid groups and stacking membranes into multilayers [28, 29]. P2 may function as a lipid transporter through a collision transfer mechanism, and it can bind cholesterol and fatty acids [27, 30–32]. Similarly to P0, mutations in P2 have been associated with the demyelinating type of CMT (CMT type 1) [33–35].

P0 spans the membrane and has domains occupying both the IPL and the MLD of compact myelin, being adhesive on both sides of the lipid bilayer. Accordingly, a common denominator for P0ct, P2, and MBP is that they all reside within the MDL of PNS compact myelin. Both P0ct [7, 10, 11, 36–42], MBP [20, 23, 37, 38, 43–48], and P2 [26, 27, 32, 33, 35, 49–51] have been studied separately with respect to their structure and membrane interactions, while their effects on lipid membranes together have received little attention. A few studies have pursued myelin protein-protein associations: P0 with myelin protein 22 (PMP22) [52, 53], MBP and P2 [54], PMP22 with calnexin [55], L-periaxin and dystrophin-related protein [56] as well as β4 integrin [57], and 2′,3′-cyclic nucleotide 3′-phosphodiesterase (CNPase) with actin [58, 59]. However, the interplay between myelin proteins remains poorly characterised at the molecular level.

Hundreds of proteins were identified in the PNS myelin proteome [60] using sensitive methods; however, the canonical major myelin proteins are the ones most important for the molecular assembly of the myelin multilayer. This is highlighted by the fact that several myelin proteins can induce membrane multilayer formation when reconstituted into lipid vesicles [7, 20, 49, 54, 61]. Myelinogenesis, sustainment of healthy myelin, and proper compaction of the MDL rely upon effective interactions between myelin proteins, and deviations in this process may cause demyelinating disease. Overall, little is known about the molecular mechanisms of myelin proteins working together.

We explored the PNS myelin proteins P0ct, MBP, and P2 together in a biomimetic membrane system by employing biophysical methods to investigate protein-lipid interactions. All three proteins induce spontaneous formation of highly ordered lattice systems, which are not purely lamellar, without affecting the structure of one another. Myelin proteins harbor key features that may synergistically shape the MDL of PNS myelin, while they on the other hand compete for membrane surface binding.

## Materials and methods

### Mutagenesis of MBP

To prepare a bacterial expression construct of MBP with a native N-terminal sequence, we used our earlier pTH27 [62] construct encoding for an N-terminally His_6_-tagged mouse 18.5-kDa MBP as template [20]. We generated a linear amplicon with part of the coding sequence deleted using Phusion HF (Thermo Scientific) for PCR (5′-phosphorylated forward primer: 5′-GCAAGCCAGAAGCGTCCG-3′; reverse primer: 5′-CTGAAAATAAAGATTCTCAGAGCCTGC-3′). Template DNA was digested with *Dpn*I (NEB), and the remaining linear amplicon was circularized using T4 DNA ligase (NEB). This resulted in a new construct, where the codons encoding for the last Gly residue in the N-terminal TEV digestion site (ENLYFQ*G*) and the initial Met of MBP (*M*ASQ…) were removed. The identity of the construct was verified using DNA sequencing.

### Expression and purification of P0ct, P2, and MBP

Expression and purification of human P0ct and human P2 have been described [7, 27]. The newly cloned mouse MBP was expressed and purified as described [20]. After TEV digestion, the obtained MBP had the exact sequence of endogenous mouse MBP after initiator Met cleavage, starting with an N-terminal Ala [63].

Final size exclusion chromatography (SEC) after affinity purification and tag removal was performed using a HiLoad Superdex 75 pg 16/60 or Superdex 75 10/300GL column (GE Healthcare) with 20 mM HEPES, 150 mM NaCl, pH 7.5 as the running buffer. Absorbance at 280 nm was used before each experiment to measure accurate protein concentrations.

### Expression and purification of MBP-C1-His and MBP-C8-His

pET22b plasmids containing either MBP-C1 or MBP-C8 [64, 65], both containing an uncleavable C-terminal His_6_-tag, were transformed into *Escherichia coli* BL21(DE3)pLysS RARE cells and selected using 100 µg/ml ampicillin and 34 µg/ml chloramphenicol. MBP-C1-His and MBP-C8-His were expressed and purified as described [20]. The final SEC buffer was 20 mM Tris-HCl, 200 mM NaCl, pH 8.0.

### Lipid preparation

DMPC (dimyristoyl phosphatidylcholine) and DMPG (dimyristoyl phosphatidylglycerol) were acquired from Larodan Fine Chemicals AB (Malmö, Sweden), and DOPC (dioleoyl phosphatidylcholine) and DOPS (dioleoyl phosphatidylserine) were purchased from Avanti Polar Lipids (Alabaster, Alabama, USA). 10 mg/ml stocks of lipid mixtures (DMPC:DMPG (1:1)) were prepared by dissolving DMPC in chloroform and DMPG in chloroform:methanol (75:25) (hyper grade for LC-MS, Merck). The solvent was evaporated under a gentle stream of N_2_ before freeze-drying the lipid film overnight at -52 °C under vacuum. The dried lipid stocks were stored at -20 °C until use.

Right before experiments, the appropriate lipid model – small unilamellar vesicles (SUVs/liposomes), large unilamellar vesicles (LUVs), multilamellar vesicles (MLVs) or bicelles – was prepared by hydration into either water (Milli-Q) or HBS (20 mM HEPES, 150 mM NaCl, pH 7.5) buffer to a final concentration of 5-10 mg/ml. Hydration was performed by continuously inverting the glass vials overnight at ambient temperature. Hydrated lipids were then subjected to seven cycles of freeze-thawing and vortexing. The formed MVLs were either used as such for DSC, or subjected to further preparation to make SUVs, LUVs, and bicelles. SUVs were made by sonicating the MLVs using a probe tip sonicator (Branson Model 450, Sonics & Materials Inc. Vibra-cell VC-130) until the lipid suspension was clear while avoiding overheating. 100-nm LUVs were prepared by extruding MLVs 11 times through a 0.1-μm membrane filter on a +40 °C heat block and used immediately. Bicelles were created by adding *n*-dodecylphosphocholine (DPC) to the MLVs to a q-ratio of 2.8 and incubating at room temperature for a couple of hours to equilibrate before use.

For membrane patch experiments, DOPC and DOPS were dissolved in methanol to a final concentration of 10 mM and a PC:PS ratio of 9:1. 0.05% DiD-C18 probe (Thermo-Invitrogen) was added to the mixture after the lipids were completely dissolved, and the lipid films were prepared immediately thereafter.

### Differential scanning calorimetry

3.5 μM P0ct, 1.75 μM P2, and 1.4 μM MBP were mixed with MLVs in HBS, at 350 μM DMPC:DMPG (1:1), in a final volume of 700 μl. This resulted in protein-to-lipid (P/L) ratios of 1:100 for P0ct, 1:200 for P2, and 1:250 for MBP, also in the protein mixtures. Lipid samples without protein were run as controls. The samples were degassed at +10 °C for 15 min under vacuum. DSC was performed using a MicroCal VP-DSC instrument, with a cell volume of 500 μl. HBS buffer was used as a reference, and the scan rate was +60 °C/h with a 4-s filtering period. Each calorimetric cycle (one per sample) was performed from +10 to +40 to 10 °C with 1 °C/min increments. Baselines were corrected and zeroed at +20 °C to facilitate cross-comparison. Calorimetric enthalpy of the phase transition (ΔH_cal_) and the apparent phase transition temperature (T_m_) were determined by integrating the DSC peak in GraphPad Prism 10. The y-value of the peak maximum was used to calculate peak width at half-maximum (ΔT_1/2_).

### Synchrotron radiation circular dichroism spectroscopy

P0ct, P2, and MBP were dialyzed into double-distilled water prior to SRCD measurements to remove salt and HEPES. The three proteins were studied individually and in protein mixtures in a 100-μm pathlength closed cylindrical cell (Suprasil, Hellma Analytics) at concentrations 50 μM for P0ct, 25 μM for P2 and 20 μM for MBP. SUVs or bicelles were hydrated in water and prepared as described above. Proteins were mixed with SUVs and bicelles right before measurements to a final P/L ratio of 1:100 for P0ct, 1:200 for P2 and 1:250 for MBP. SRCD data were collected on the AU-CD beamline at the ASTRID2 synchrotron (ISA, Aarhus, Denmark). Spectra were recorded from 170 to 280 nm at +30 °C, with 6 scans per measurement. The baseline was subtracted before comparing the CD spectra.

### Turbidimetry

For turbidimetry, 5 μM P0ct, 2.5 μM P2, and 2.0 μM MBP were measured individually or in various mixtures with either 0.5 mM DMPC:DMPG (1:1) liposomes or 0.5 mM DMPC:DMPG (1:1) bicelles in HBS. All samples were measured in duplicate at a final volume of 100 μl on a Greiner 655161 96-well plate. Optical density at 450 nm (OD_450_) was recorded immediately after mixing at +30 °C using a Tecan Spark 20M plate reader. HBS buffer alone and 5 μM BSA were used as negative controls. Buffer was subtracted, and OD_450_ at the highest point during each measurement (t = 3 min for liposomes, t = 2 min for bicelles and t = 1 min for HBS) was then plotted for comparison. The signal deteriorated after this maximum, due to the aggregation of large proteolipid particles.

### Surface plasmon resonance

Surface plasmon resonance (SPR) was performed on a Biacore T200 system (GE Healthcare). 100-nm LUVs of 2 mM DMPC:DMPG (1:1) were immobilized on an SCBS LD sensor chip (Xantec Bioanalytics) in HBS, followed by injection of P0ct, P2, and MBP mixtures at +30 °C. Chip regeneration was done with 40 mM (3-[(3-cholamidopropyl)dimethylammonio]-1-propanesulfonate) (CHAPS) followed by 50 mM NaOH. To investigate if the proteins affected each other upon lipid binding, 0.5 - 1 mM of each protein (P0ct, MBP, and P2) was injected after each other in various sequences upon the same lipid capture. Each response curve was zeroed at time = 2600 s (right before the first protein injection) and plotted. For each run, HBS was run three times as a control for the experiment, and all samples were run in duplicate.

For protein binding affinity and kinetic analysis, 100-2000 nM of each protein was injected with a single concentration per lipid capture. All samples were performed in duplicate, with one sample in each series measured twice to rule out instrumental artefacts or deviations. To gain information about association affinity, the binding response as a function of protein concentration was plotted and fitted to a 4-parameter model:

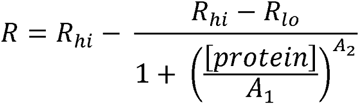

Kinetic analysis was executed as follows: all association phases (180 s after the start of protein injection) were individually fitted to a one-phase exponential association model using GraphPad Prism 10. Protein concentration was plotted against the obtained *k_obs_* values, and *k_on_* (slope of the curve) and *k_off_* (Y-intercept of the curve) were determined by using linear regression. *k_on_* and *k_off_* were extracted from two fitted datasets as described [20].

### Transmission electron microscopy

P0ct, P2, and MBP were mixed with 1 mM DMCP:DMPG (1:1) SUVs or bicelles in HBS to a final P/L of 1:100, 1:200, and 1:250, respectively, before incubating for 1 h at room temperature. The final samples were incubated for 1 h before 4 μl of each were added onto a carbon-coated copper grid, 200 mesh (lines/inch), and incubated for 1 h. Excess liquid was then carefully removed with Whatman filter paper. Each grid was stained with 2% uranyl acetate for 12 s before the staining solution was removed and the samples were left to air dry. TEM imaging was performed using a Hitachi HT7800 TEM instrument. The samples were prepared in duplicate, and images were taken at several locations on each grid. Images were finalized in ImageJ [66].

### Small-angle X-ray diffraction

SAXD experiments were performed to investigate ordered, repetitive structures in proteolipid assemblies. P0ct, P2, and MBP were mixed with SUVs or bicelles of 3 mM DMPC:DMPG (1:1) in HBS at room temperature and exposed to X-rays on the CoSAXS synchrotron beamline, MAX IV Laboratory, Lund University (Lund, Sweden) [67]. To achieve P/L of 1:100, 1:200 and 1:250, 30 μM P0ct, 15 μM P2, and 12 μM MBP were used, respectively. A combined concentration of 50 μM, resulting in a P/L of 1:60, was used as a control. Lipid samples without protein displayed the typical lipid membrane form factor, without Bragg peaks. X-ray energy was fixed at 12.4 keV (wavelength 1.0 Å). Exposure time was 200 ms, collecting 300 frames per sample. Data were processed and analysed using ATSAS [68].

The diffraction data are displayed as intensity *I*(*s*) as a function of modulus of the scattering vector *s*:

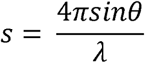

2θ being the scattering angle. The Bragg peaks were indexed to identify possible lipid liquid crystalline phases present in the samples. A lamellar phase has Bragg peaks at *s* = *h* 2π/*a,* where *a* is the repeat distance and *h* is an integer (Miller index), and a hexagonal phase has Bragg peaks at 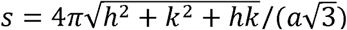, where *a* is the unit cell side length and *h* and *k* are integers (Miller indices). For the cubic phases, the Bragg peaks are at 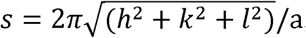, where *h*, *k*, and *l* are the Miller indices with integer values and side length of the cubic unit cell. Depending on the symmetry of the cubic phases, some diffraction peaks are allowed while some are forbidden [69–71]. By indexing the diffraction peaks one can determine the symmetry of the cubic phases. Usually, for lipid systems, four different types of cubic phases are observed, which are Im3m, Ia3d, Fd3m or Pn3m. For the samples in the present study, we have found that in presence of liposomes, two different types of self-assembled structures were formed; lamellar, Im3m or Fd3m, whereas in presence of bicelles, cubic phases of Ia3d and Im3m symmetries are observed. Representative examples for lamellar, Im3m/Fd3m, Ia3d and Im3m in Supplementary Figure 3 and Supplementary Table 3.

### Time-lapse epi-fluorescent microscopy

Hydrated double-supported lipid membrane patches were prepared as previously described [72, 73]. To prepare planar lipid bilayers, 50 μl of 10 mM DOPC:DOPS (9:1:0.05%DiD-C18) was applied to a freshly cleaved Mica substrate (Plano GmbH), which was glued to a glass coverslip using silicone elastomer (MED-6215, Nusil Technology). Excess lipid solution was removed by spinning the mica on a spin coater (KW-4A, Chemat Technology) at 150 g for 40 s. Solvents were evaporated under vacuum in a desiccator for 10-12 h. The lipid films were hydrated in 10 mM Tris-HCl, 140 mM NaCl, 2 mM Ca^2+^, pH 7.4, for ≥1 h at +60 °C. Defined secondary bilayer patches were prepared by buffer-exchange >10 times, before cooling the patches down to room temperature and letting them equilibrate for >1 h before experiments.

Addition of myelin proteins and the response of the subjected bilayer patches was monitored with time-lapse *epi*-fluorescence microscopy using a Nikon Ti2-E inverted microscope with a 40× objective (Nikon ELWD S Plan Fluor, NA = 0.6). Fluorescence excitation at 550 nm was done with a white light fluorescence illumination system (CoolLED, pE-300), and the emission at 640 nm was monitored (Nikon Cy5-A, M376571). Images were captured at 10 fps with a digital CMOS camera (ORCA-Flash 4.0 V3, Hamamatsu, 2048 × 2044 pixels) and NIS-Elements software (Nikon).

For the samples containing 10 μM P0ct, 10 μM MBP-C1-His and 10 μM MBP-C8-His, 5 experiments were run per condition. Preliminary testing of various protein combinations (Supplementary Figure 8, Supplementary Movies 4-8) was run at 2.77 - 20 μM protein, depending on the protein(s) in question. Limited amounts of P0ct, MBP, and P2 restricted the number of tests in these studies. These experiments were only executed once, and further testing will be needed to verify the findings. Before experiments, P0ct, MBP-C1-His and MBP-C8-His were dialysed into water, snap-frozen with liquid N_2_ and freeze-dried at -52 °C before rehydrating them in 10 mM Tris, 140 mM NaCl, 2 mM Ca^2+^, pH 7.4. For preliminary tests on protein mixtures, P0ct, MBP-C1 and P2 were dialyzed directly into 10 mM Tris, 140 mM NaCl, 2 mM Ca^2+^, pH 7.4.

## Results

P0, MBP, and P2 have been extensively studied as individual proteins [5, 7, 9–11, 20, 21, 35, 37, 38, 49, 51, 52, 74–77], but a more comprehensive study of them functioning together in interacting and stacking lipid membranes in a myelin sheath model has been lacking. We set out to test the effects of mixing these three proteins, using biophysical methods to follow potential synergy, competition, bilayer stacking, membrane binding and other effects. To mimic P/L ratios found within the native PNS myelin sheath, and to consider requirements of *in vitro* studies of the protein-lipid complexes, we used P/L ratios 1:100 for P0ct, 1:200 for P2, and 1:250 for MBP. These three P/L ratios were used in all experiments unless otherwise stated.

### The combination of myelin proteins prompts a large reduction in lipid phase transition temperature

Lipid phase transition temperatures reflecting the conformational freedom of the lipid hydrocarbon tails were explored using DSC. The proteins were screened against DMCP:DMPG (1:1) MLVs, monitoring any protein-induced effects on the endothermic transition (T_m_) of the lipids (Figure 1). The phase transition of the lipid system from gel (L_β_) to fluid (L_α_) phase can be investigated based on the DSC thermogram together with potential pre-transition phases, such as rippled gel (P_β’_) and inverted hexagonal (H_II_) phases [78].

**Figure 1:**
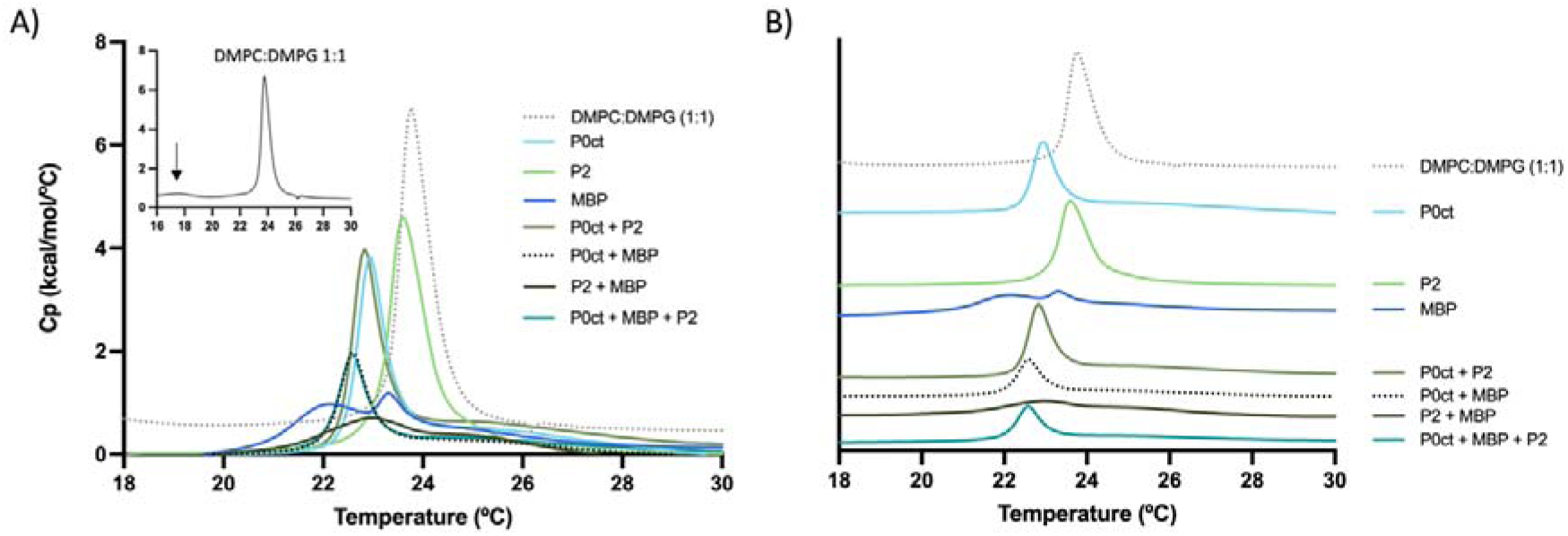
DSC measurements of MLV-protein mixtures. A) The MLVs were measured alone as control (dotted grey line, black solid line in insert). Pre-transition of the DMPC:DMPG MLVs is indicated with an arrow in the insert. The three proteins were measured at P/L 1:100 for P0ct, P/L 1:200 for P2, and P/L 1:250 for MBP. B) Same graphs as in A, moved relative to each other for clarity. Note how the phase transition occurs at lower temperatures with all protein-containing samples. The cooling scans are shown in Supplementary Figure 1.

The MLVs displayed a pre-transition at +17.5 °C (Figure 1A, arrow) and an endothermic transition at +23.70 °C, similarly to earlier findings [11, 79, 80]. However, the addition of all three myelin proteins eradicated the pre-transition peak and decreased T_m_, thus altering the lipid phase transition behavior. The addition of protein decreased the main transition population with an appearance of a shoulder at ∼ 25.0 °C in several samples. MBP at P/L 1:250 was the only protein that induced two phase transitions, displaying a broader peak at +22.15 °C and a second peak at +23.30 °C. Besides MBP, the addition of the three proteins together had the most effect on the melting temperature, reducing the T_m_ by 1.21 °C. From each DSC thermogram, the area under the peak was calculated, from which the calorimetric enthalpy of the transition (ΔH_cal_) and temperature width at half-height (ΔT_1/2_) could be estimated (Table 1). The enthalpy of the phase transition was notably reduced with the addition of protein(s), whereby P2 + MBP nearly eliminated any detectable (L_β_) - (L_α_) phase transition. The cooperativity of the system was examined based on ΔT_1/2_, and P0ct was the only protein displaying properties of stabilizing the MLVs phase transition, suggested by the reduced ΔT_1/2_.

**Table 1:** DSC results. The phase transition temperature (T_m_), change in T_m_ (ΔT_m_), calorimetric enthalpy of the transition (ΔH_cal_), and temperature width of half-height (ΔT_1/2_) of the DSC heating trace from the various protein mixtures compared to the MLVs alone. Additionally, ΔT_m(heating-cooling)_ is shown for DSC heating and cooling scans. Cooling scans are shown in Supplementary Figure 1.

| Mixture | $T_m$ (°C)<br>Heating | $\Delta T_m$ (°C)<br>Heating | $\Delta H_{cal}$<br>(Kcal/mol) | $\Delta T_{1/2}$<br>(°C) | $\Delta T_m$ (°C)<br>Heating - Cooling |
| --- | --- | --- | --- | --- | --- |
| DMPC:DMPG 1:1 | 23.77 | - | 16.4 | 0.787 | -1.43 |
| P0ct | 22.96 | -0.81 | 4.98 | 0.650 | -2.26 |
| P2 | 23.57 | -0.20 | 5.71 | 0.814 | -1.06 |
| MBP | 22.15, 23.3 | -1.62, -0.47 | (4.54) | (2.840) | -1.75, -0.80 |
| P0ct + MBP | 22.56 | -1.21 | 2.68 | 0.697 | -2.30 |
| P0ct + P2 | 22.83 | -0.94 | 6.30 | 0.681 | -2.15 |
| P2 + MBP | 22.96 | -0.81 | 2.30 | 2.683 | -2.69 |
| P0ct + P2 + MBP | 22.56 | -1.21 | 3.14 | 0.700 | -2.32 |

Analogous types of phase transitions were generated when the mixtures were cooled down again (Supplementary Figure 1A), but the reverse L_α_ - L_β_ phase transitions shifted to a lower temperature range (Supplementary Figure 1B). The T_m_ for MLV conversion to the gel phase was +22.43 °C, a difference of -1.43 °C from the heating scans (Table 1). As individual proteins, both MBP and P2 had a slight positive effect, with a +0.07 °C and +0.08 °C increase of the T_m_. Again, P0ct, P2, and MBP together had the greatest effect of -2.16 °C.

### Little impact on secondary structure between proteins

SRCD was utilized to study any protein-protein interaction-induced conformational changes in the proteins in the presence of lipids. The three proteins were measured individually (Supplementary Figure 2) and as mixtures (Figure 2), and the mixtures were compared to calculated combined spectra from individual proteins. As previously shown, P0ct and MBP remained unfolded in solution and gained α-helical content when added to lipidic systems [7, 11, 20, 37], while P2 displayed a β-barrel conformation (Supplementary Figure 2) [26, 35, 49, 51]. Overall, the secondary structure of each protein was not affected by the other proteins, nor when the lipidic model system changed from liposomes to bicelles. The exception was the P2 + MBP sample (Figure 2C); a small spectral shift is noticeable in the bicelle sample, as the ellipticity peaks move 2 nm between calculated and measured spectra. This can indicate either that the P2 and MBP alter their secondary structure or that they together cause slight alterations in the protein-lipid environment, distinct from when the individual proteins are measured alone [81]. Another option is that they compete for binding, and in the mixture, not all P2 or MBP is bound onto the membrane in the sample.

**Figure 2:**
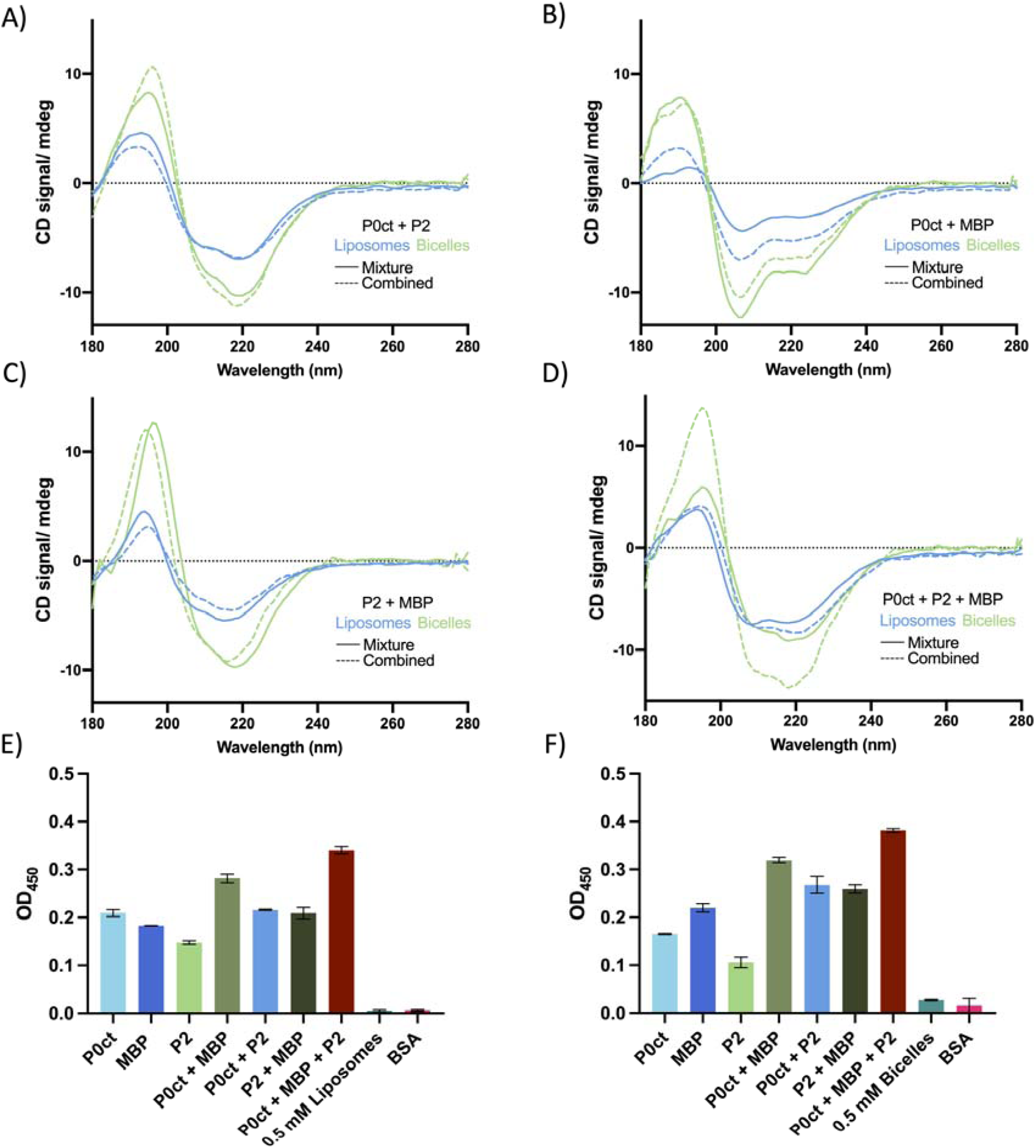
CD spectroscopy and turbidimetry. SRCD measurements of protein mixtures added to either liposomes (blue) or bicelles (green), plotted with “combined” spectra (dotted lines), which show the sum of each protein component measured individually. In Supplementary Figure 2, the SRCD spectra of each individual protein are shown measured in solution, liposomes. and bicelles. The following mixtures are shown: A) P0ct + P2, P/L 1:100 and 1:200. B) P0ct + MBP, P/L 1:100 and 1:250. C) P2 + MBP, P/L 1:200 and 1:250 and D) P0ct + P2 + MBP, P/L 1:100, 1:200, and 1:250, respectively. Each spectrum was measured at the wavelength from 280 – 170 nm performing 6 scans per sample at +30 °C. E-F) Liposome and bicelle turbidimetry. Induced turbidity of P0ct, MBP, and P2, or mixtures thereof, were studied in DMPC:DMPG (1:1) liposomes and bicelles at constant P/L ratios of 1:100 for P0ct, 1:200 for P2, and 1:250 for MBP. 5 μM BSA (P/L 1:100) was used as a negative control. OD_450_ was measured for 20 min at +30 °C (Supplementary Figure 9). Each condition in the absence of liposomes/bicelles is shown in Supplementary Figure 9.

### Synergistic effects on protein-induced turbidity

The aggregation behavior of the individual myelin proteins has been investigated using vesicles [7, 11, 20, 26, 37], and the turbidity induced by each individual protein supports earlier findings (Figure 2E-F, Supplementary Figure 9). The protein combinations boosted the measured turbidity for both membrane models, indicating synergistic effects between the proteins. Turbidity was essentially absent in control samples (Supplementary Figure 9C-D). In all seven myelin protein samples, the greatest effect on turbidity was seen when all three proteins were present, with slightly higher turbidity for bicelles compared to liposomes (Figure 2E-F). These data are semi-quantitative, due to the chemical differences in the lipid systems, as well as the measurement time point being subjectively taken at the peak position before bulk aggregation starts to decrease the signal.

### Membrane surface binding

Binding kinetics and affinity of P0ct, P2, and MBP towards membrane surfaces have been investigated using SPR in earlier work, revealing different kinetic properties for each protein [7, 20, 51]. We wanted to explore the association to vesicles when the proteins were injected as mixtures and when they were added in a defined sequence, to shed light on potential cooperation and/or competition between the three myelin MDL proteins (Figure 3). Binding kinetics were studied for each individual protein (Supplementary Figure 4), and similarly to earlier findings [7, 20, 51], MBP had the highest responses (R_hi_) for DMPC:DMPG 1:1 vesicles and P2 had the weakest R_hi_ towards the LUVs (Supplementary Table 1). MBP and P0ct bound irreversibly to the immobilized vesicles, whereas most P2 dissociated rapidly (Figure 3A). The obtained saturation midpoint concentrations (A_1_), or apparent dissociation constants (K_d_), were in a comparable range (0.5-1 μM) to earlier studies for all three proteins (Supplementary Figure 4; Supplementary Table 1-2) [7, 20, 51].

**Figure 3:**
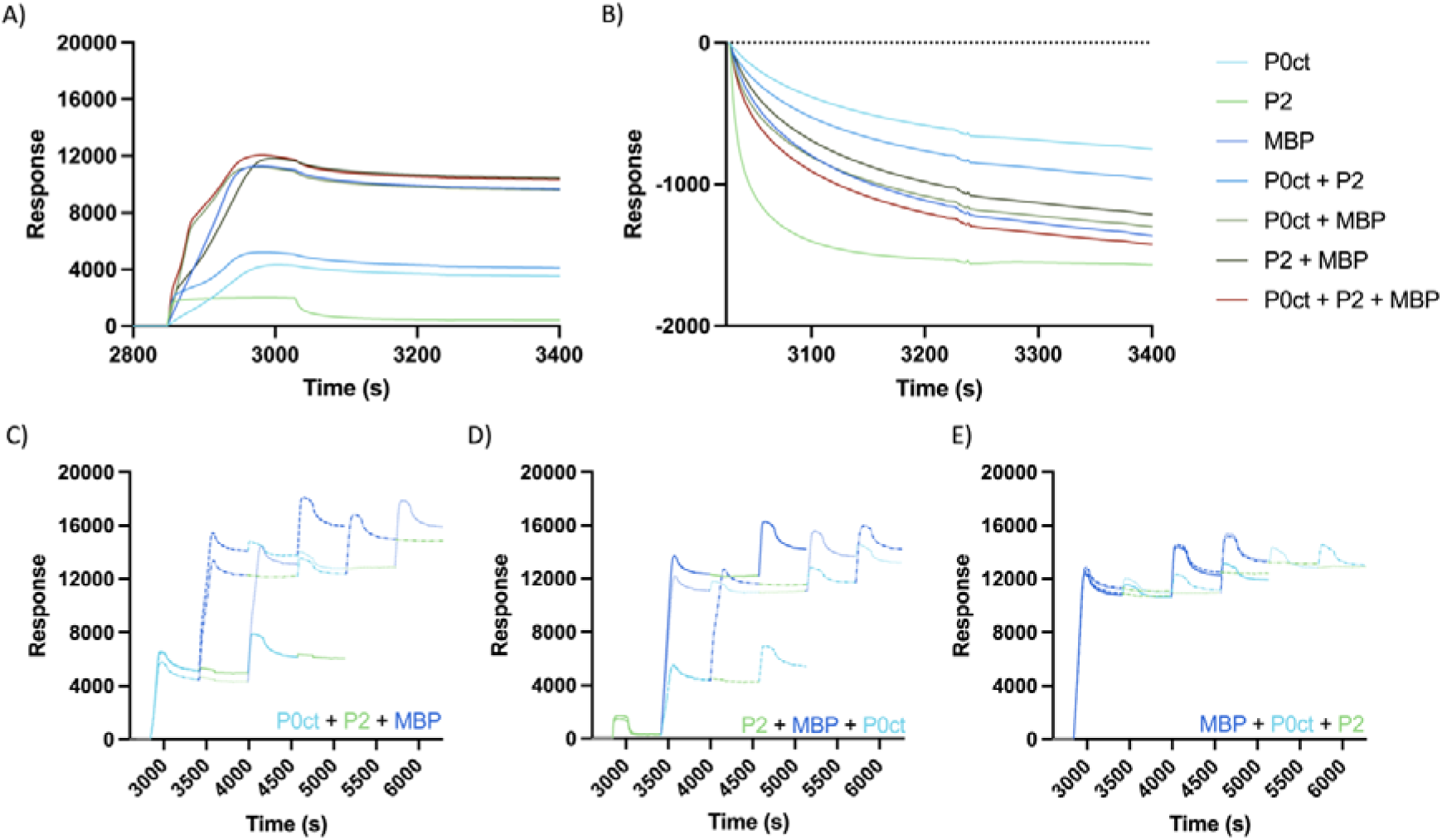
SPR analysis of membrane interactions. A) Measurements of P0ct, P2, and MBP injected as protein mixtures. B) Dissociation from immobilized lipid vesicles. Zeroed at t = 3027 s. C-E) Proteins injected in sequence, where each protein was set to be the “start” protein in the sequence. P0ct – light blue, P2 – green, MBP – dark blue. The concentration for each protein was 1 μM. Supplementary Figure 6 shows each protein sequence separately.

Protein association with immobilized vesicles was first investigated by adding the three proteins in various mixtures (Figure 3A), and the shape of the association phase depended on which proteins were present. P2 generated a rapid increase in response, while P0ct bound to the membranes slower, as indicated by the prolonged association phase. Furthermore, the highest response was obtained with a combination of the three proteins, as well as with P2 and MBP. Dissociation of the proteins from the vesicles was investigated by zeroing the data at the end of protein injection (after 180 s), as viewed in Figure 3B. Interestingly, P2 appeared to be dependent on binding partners to not dissociate from immobilized vesicles. Increased amounts of P2 bound irreversibly to the lipid surface when P0ct or MBP were present (Figure 3B). Out of the four protein mixtures, the combination of P0ct, P2, and MBP dissociated more than any other mixture, displayed by the decrease in response, potentially due to membrane surface saturation and protein competition.

Competition between the proteins was investigated by injecting them in all possible sequential orders and visually interpreting the binding to the immobilized vesicles (Figure 3C). From the first injection in each setup, it could be seen that P2 bound at 2000 RU, P0ct at 6-7000 RU, and MBP at 12000 RU onto a protein-free membrane under the employed conditions. While P2 dissociated rapidly and nearly completely, MBP and P0ct remained mainly on the membrane and inhibited the subsequent binding of each protein, indicating both membrane saturation and competition between the proteins. When P0ct was added as the initial protein, MBP still bound strongly to the lipid vesicles, as seen with the increase in RU (Figure 3C). In contrast, P2 only managed to bind in small amounts (∼300 RU, Figure 3A) or not at all. This was also the case when either MBP or P2 was injected as initial proteins (Figure 3D-E). Most P2 dissociated from the immobilized vesicles rapidly, but a permanent increase was seen at roughly ∼ 400 RU (Figure 3D), establishing a minor quantity of P2 attached stably to the membrane before it was exposed to MBP or P0ct. Decreasing the concentration of P0ct and MBP to 0.5 μM (Supplementary Figure 5) did not increase the P2 association. The results show an interplay between the three proteins, whereby there is a limit to the amount of bound protein on the membrane surface, and the presence of bound myelin protein on the membrane inhibits the binding of additional protein.

### From bilayer structures to evidence of hybrid lipid crystals

X-ray diffraction has been used to investigate the membrane stacking function of P0ct, MBP, and P2 [7, 20, 39, 49]. Bicelles have shown enhanced stacking order and reveal latent protein-induced phase transitions beyond the usual gel (L_β_)-to-liquid crystalline (L_α_) phase [73]. Liposomes and bicelles do not show Bragg peaks on their own, and the usual form factor is displayed from each of the lipid models, similarly to what has been published before [73]. When subjected to liposomes, individual proteins in native-like concentrations induced a Bragg peak corresponding to a lamellar d-spacing ranging between 74-90 Å (Figure 4A,C, Supplementary Table 3). Wider distances were only seen when P2 was measured alone, and P0ct and MBP both caused the repeat to shorten to ∼ 73-76 Å regardless of whether P2 was present in the sample. Higher concentrations of MBP induced two weaker peaks at around s = 0.11 and 0.14 Å^-1^ (Figure 4C) that have been observed before [26]; these Bragg peaks are not related to the lamellar d-spacing but matched the lattice parameters of an Im3m/Fd3m cubic mixture (Supplementary Figure 3). Similarly, several of the control protein-liposome samples indicated a formation of Im3m/Fd3m cubic phases at the cost of the lamellar phase.

**Figure 4:**
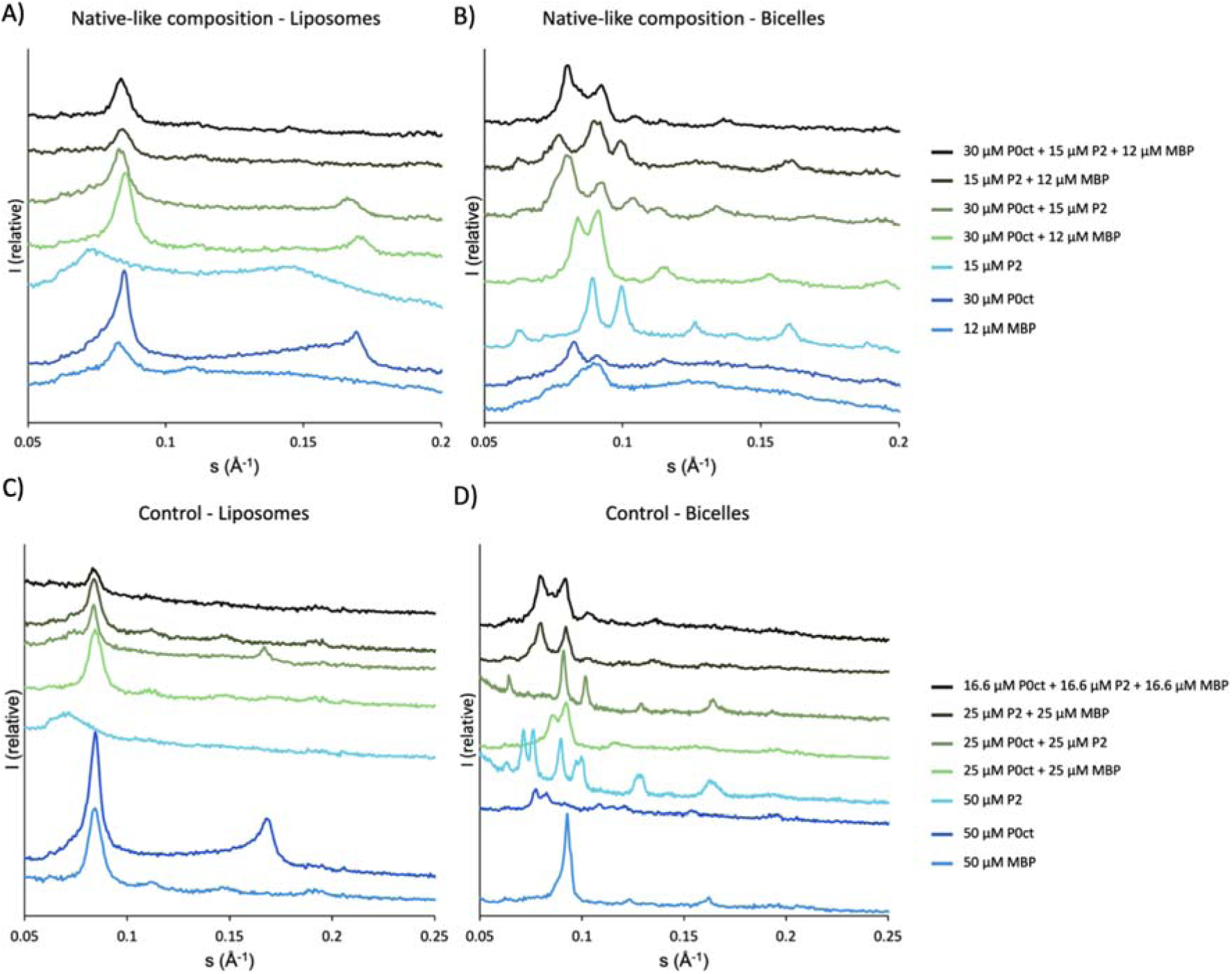
Mixing of the P0ct, P2, and MBP individually and in various mixtures with DMPC:DMPG (1:1) SUVs and bicelles results in X-ray diffraction patterns with several Bragg peaks. A-B) “Native” protein-to-lipid ratios (P/L) of proteins referred to as “samples” whereas C-D) “control” mixtures with an unusually high protein concentration were mixed with 3 mM SUVs/bicelles. The spectra have been offset for clarity and the data have not been scaled to one another. Possible lattice systems and indexing are shown in Supplementary Figure 3 and Supplementary Table 3.

When bicelles were used, up to eight diffraction peaks were induced depending on the protein composition (Figure 4B,D), and additional peaks emerged at higher protein concentrations (Figure 4D). Diffraction analysis revealed that the native-like protein compositions induced lattice parameters with symmetries fitting cubic phases of Im3m or Ia3d (Supplementary Figure 3, Supplementary Table 3), while each protein alone suited a mixture of Ia3d/Pn3m or Im3m/Ia3d.

### Bicelle aggregates develop into filament-like structures in protein mixtures

The samples resembling native-like P/L ratios were visualized with TEM. As shown in Supplementary Figure 7, individual proteins induced typical protein-bicelle aggregates [39, 49]. Interestingly, the different mixtures showed modifications to the standard cluster morphology, displaying both longer and shorter strand-like structures (10 μM P0ct + 4 μM MBP) and extensive bundles of apparent filaments or tubes of fused bicelles (10 μM P0ct + 5 μM P2 + 4 μM MBP) (Figure 5). The sample of P0ct + P2 stood out with cleaner grids, where all lipid in the sample appears to be concentrated into the aggregates, whereas the strands in the P0ct + MBP blend were scattered across the whole grid, being visible even at low magnifications. Even though there were clusters in the P2 + MBP sample, several filamentous constellations were identified across the grids (Figure 5). Similar structures can be observed when all three proteins are present.

**Figure 5:**
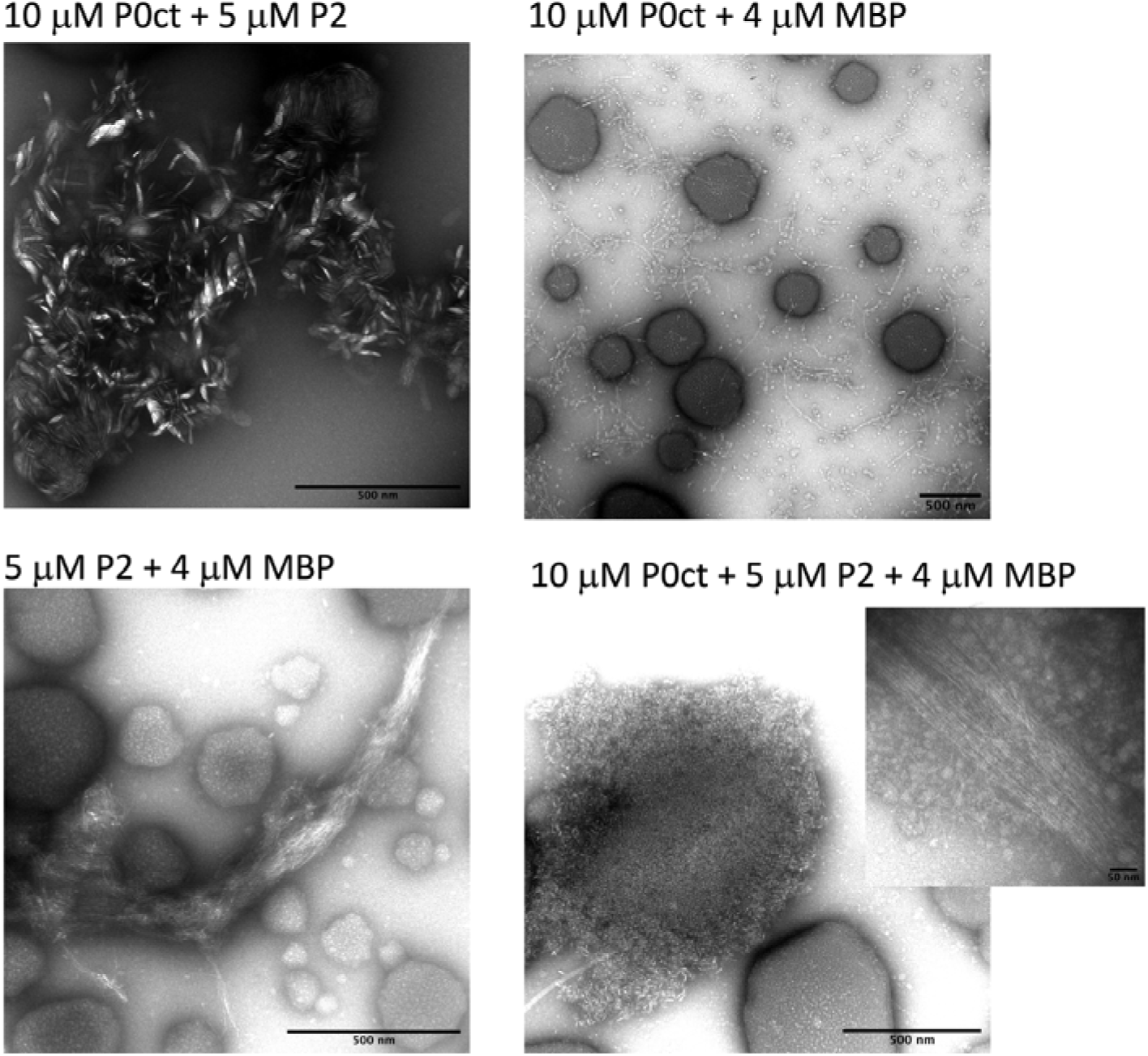
TEM images of mixed proteins. 1 mM bicelles (DMPC:DMPG 1:1) were mixed with 10 μM P0ct, 5 μM P2, and 4 μM MBP, to match protein-to-lipid ratios of 1:100, 1:200, and 1:250, respectively. The addition of individual proteins to 1 mM bicelles can be viewed in Supplementary Figure 7.

### Myelin proteins P0ct and MBP induce the formation of new membrane structures and membrane patch expansion

To study the possible effects of myelin proteins (P0ct, MBP-C1-His and MBP-C8-His) on membranes, planar double supported membrane patches having the structure shown in Figure 6 (top) were utilized. After addition, the proteins reached the membranes within 30-60 s.

**Figure 6:**
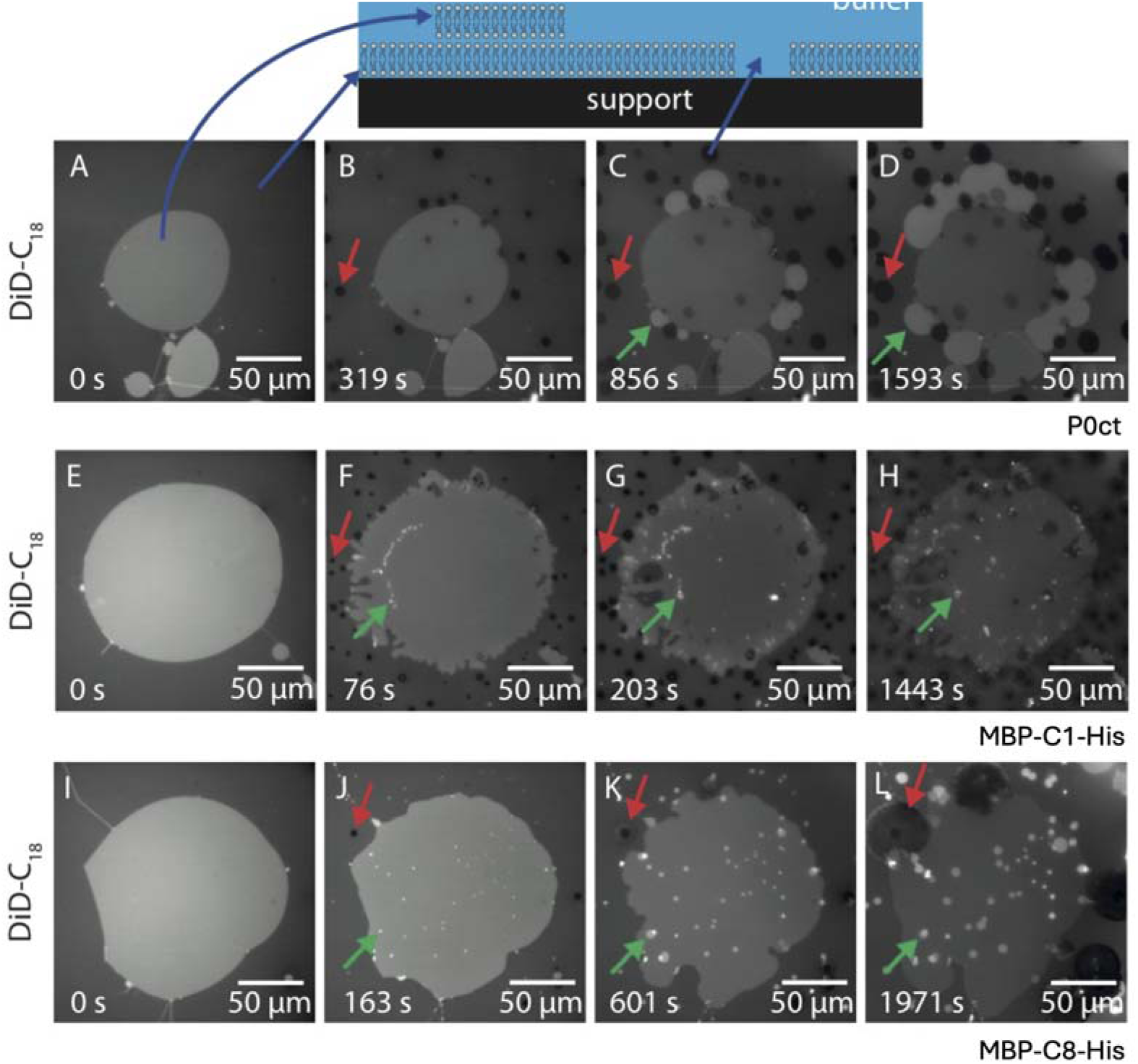
Fluorescence patch experiment of A-D) P0ct and of the two isoforms E-F) MBP-C1-His and I-L) MBP-C8-His. 10 μM of each protein was added to DOPC:DOPS (9:1, 0.05% DiD-C18) bilayers. Before adding protein (0 s), the patches and the lipid bilayer. P0ct, MBP-C1 and MBP-C8 induce cavities in the underlying bilayer (red arrows). The formation of new membrane bilayers is indicated with green arrows. The effects of protein addition were visualized for 15 min. Original movies are shown in Supplementary Movies 1-3. The fluorescent bilayers become bleached with time due to the prolonged light exposure. An illustration of the double-supported membrane structure can be viewed at the top.

Fluorescence patch experiments of P2 and P0ct have been published earlier [35, 73], whereby P0ct induced cavities and lipid accumulation potentially due to membrane condensation, whereas P2 primarily prompted new bilayer structure formation without affecting underlying membranes. Here, the addition of myelin proteins P0ct, MBP-C1-His and MBP-C8-His resulted in the formation of new membrane structures, and the creation of holes in the supporting membrane as visualized by time-lapse fluorescence microscopy (Figure 6). First, the small holes appeared in the supporting membrane, and then, new membrane structures emerged from the double supported membrane patch; both the holes and the new membrane structures grew over time. For P0ct, the new membrane structures created were larger relative to the smaller structures induced by MBP-C1-His and MBP-C8-His. The proteins reached the membranes within 30-60 s, with smaller membrane structures materializing around the edges of the existing bilayer. The membrane structures induced by wild-type P2 [35] were of similar size compared to the observation for P0ct in Figure 6A-D, whereas the P2 mutants induced smaller membrane structures, which resemble the MBP-induced effects in Figure 6E-L. MBP, on the other hand, appears to induce disorder on the bilayer edges, strongly shrinking the original double-supported membrane patch (Figure 6E-H), while tiny lipid structures emerge on top of it. Cavities, same as for P0ct, can be seen in the primary membrane layer. The less cationic isoform of MBP (MBP-C8) induced swelling of the supported lipid bilayer instead of the condensation seen with MBP-C1 (Figure 6I-L).

Preliminary testing of several protein mixes was performed (Supplementary Figure 8, Supplementary Movies 4-8) with various concentrations of the proteins to acquire initial results on their cooperative effects on supported bilayers. At 10 μM MBP-C1, inflation of the double-supported membrane bilayer was observed (Supplementary Figure 8A-E), similarly to the effects of high concentrations of the C8 isoform. At even lower concentrations, MBP-C1 can induce bilayer formation together with P0ct (supplementary figure 8F-J). Equal concentrations of P0ct and P2 generated only a small growth of new bilayers, whereas doubling the P2 concentration produced a larger membrane structure kept within the margins of the original one (Supplementary Figure 8K-T). Increasing the concentration of P0ct led to both swelling and development of secondary and tertiary bilayer formation, indicated by the formation of brighter bilayer formations (Supplementary Figure 8U-Y). The preliminary findings do suggest that the combined effects of these proteins stimulate new membrane formation and elongation of existing bilayers. These results need to be further investigated in detail to verify the findings described here.

## Discussion

Maintaining the narrow apposition of the major dense line relies upon accurate combined forces from several myelin proteins. In the PNS, one glycoprotein and two basic proteins, *i.e.* P0, P2, and MBP facilitate and aid in keeping the lipid bilayers tightly together [7, 8, 10, 20, 26, 49, 82]. Here, we used several methods to study protein-lipid interactions, to obtain information on how myelin proteins cooperate at the molecular level to accomplish dense packing of the membranes at the MDL. The lipid models used were simplistic, aiming at having well-behaved lipid particles through a line of experiments, while having a negative surface charge on the membrane. Studies on more native-like lipid compositions on model membranes will shed additional light on myelin protein-lipid complex structure and cooperation.

Addition of several myelin proteins to MLVs of DMPC:DMPG (1:1) had the largest effect on the lipid phase transition temperature (T_m_), and all proteins decreased the measured enthalpy (Table 1), showing that P0ct, P2, and MBP in combination disrupt the membrane structure, lessening the van der Waals forces between the acyl chains, causing an order-to-disorder shift [83]. Decrease in ΔH_cal_ in a concentration-depended manner for P0ct, P2, and MBP has previously been observed [7, 20, 51]; each protein binds to the lipids with both electrostatic forces and hydrophobic interactions, with partial penetration into the core of the membrane system [78]. Accordingly, the enthalpy diminishes even more, when several proteins are added at the same time. Interestingly, P2 + MBP has the same impact on the T_m_ as P0ct, but the peak is almost gone. Consequently, the MLVs have barely any phase transition between the gel and fluid phase [84]. The shape of each DSC peak can reveal the miscibility between lipids and proteins, and an ideal mix between L_β_ and fluid L_α_ phase can be determined by the shape of the measured peak [85]. P0ct and MBP do not change the phase transition of pure DMPC lipids [7, 20], but most of the protein mixtures generate a broad shoulder on the right side of the main transition peak of the MLVs, which can be attributed to electrostatic interactions between the positively charged proteins and the negatively charged DMPG lipid and stabilization of the gel phase [80, 86]. On the other hand, the addition of MBP causes the peak to transform into a double peak, with a wide peak at ∼ +22 °C, demonstrating that the protein may favor one lipid component over the other. Thus, MBP binds favorably to DMPG, separating the lipids with domains enriched in DMPC, and stabilizing the fluid phase [86, 87]. MBP can phase separate in solution, creating an MBP meshwork driving protein segregation in myelin, and it has been suggested that this feature can affect lipid membranes [82]. Thus, the separation of DMPG and DMPC lipids further supports the notion that MBP associates with lipid microdomains in myelin [88]. The parameter ΔT_1/2_ can indicate relative cooperativity of the phase transition, whereby low ΔT_1/2_ denotes high cooperativity. All samples containing P0ct led to a narrowing of the phase transition, decreasing ΔT_1/2_, while P2 and MBP both gave rise to a broader peak (Table 1). While both MBP and P2 seemed to weaken the lipid membrane, the addition of P0ct caused a stabilizing effect when the proteins were combined. Hence, the overall consequence might be that P2 and MBP alter the fluidity of the phospholipids, inducing some form of intermediate states, whilst P0ct counteracts this effect, promoting ordered lipid packing. Together, these interactions lead to charge neutralization of the bilayers and membrane stacking into myelin-like structures [83, 84, 89].

Upon cooling, the L_α_ - L_β_ phase transitions are reversible while the pre-transition peak remains gone. Even though the peaks of each sample are analogous to the heating endotherm, the conversion from L_α_ phase to L_β_ phase shows cooling hysteresis (Supplementary Figure 1B), with the most profound hysteresis in the P2 + MBP sample reaching T_m_ at 2.69 °C below the heating endotherm. Interestingly, MLVs mixed with either P2 or MBP displayed peaks with a minor increase in T_m_ almost identical to the exotherm of MLVs. This may suggest that P2 dissociates completely from the MLVs after heating, as it does with immobilized vesicles in SPR. MBP, on the other hand, showed two peaks, indicating that the vesicles have protein-free lipid domains that transition to L_β_ before protein-rich domains do. Various mixtures of DMPC:DMPG exhibit minor cooling hysteresis [90], and it is clear from these results that the proteins not only affect the endotherm L_β_ - L_α_ transition, but also affect the exotherm L_α_ - L_β_ transition, emerging a stabilizing effect on the MLV liquid crystalline L_α_ phase.

For both P0ct and MBP, the lipid environment is crucial for the disorder-to-order transition of the proteins, as evident from the SRCD data (Supplementary Figure 2); P2 is folded in aqueous solution. Although combinations of the proteins have large effects on the lipid membrane properties, little protein-protein interaction is evident when investigating secondary structure changes using SRCD. A small shift in the spectra for P2 + MBP (Figure 2C) is noticeable, which is indicative of a possible contact interface between P2 and MBP, whereby one of the proteins gains more secondary structure than expected based on the individual spectra. MBP is known to have many binding partners in addition to lipids, such as calmodulin [91] and actin [92], and several studies have revealed MBP in complex with proteolipid protein (PLP) and S-MAG [93–95]. As an IDP, MBP has conformational plasticity and depending on interaction partners, it may have a diverse set of secondary structures [96]. The hydrophobic tip of P2 is postulated to be crucial for membrane binding [26], but the same domain might interact with hydrophobic residues of MBP, as shown with the IDP p53 and MDM2 [97]. Direct protein-protein interactions between the three myelin proteins studied here have not yet been reported, and further extensive studies will be required as a follow-up of the current study.

Association studies onto immobilized liposomes verified earlier findings of apparently irreversible binding of P0ct and MBP [7, 20], while P2 dissociated ∼ 6 min after the 180-s association phase is complete (Figure 3A), leaving little protein left on the membrane surface. Injecting the three proteins in various sequences clearly shows that MBP binds the strongest to the immobilized vesicles and is not much affected by the presence of P0ct or P2. When MBP or P0ct already reside on the membrane surface, similar injections of all proteins result in lower binding to the surface, indicating competition between the proteins (Figure 3C-E). Especially the binding of P2 is nearly inhibited by the presence of MBP or P0ct. The binding of MBP appeared to be more concentration-dependent than the other two proteins, since decreasing MBP concentration to 0.5 μM caused a fall in response to approximately half, while the response of P0ct remains the same (Supplementary Figure 5). Interestingly, the association of P0ct seems to weaken when a higher concentration of P2 is injected first even though most of P2 dissociates before P0ct is added. The response drops to more than half and does not reach the same values as for any other measurement. On the other hand, more P0ct bound when MBP concentration was lowered. Independently of the order of the protein injection sequence, MBP seems to be the only protein that still finds lipid domains to bind after several injections of all proteins. In the end, neither P0ct nor P2 can bind, while injections of MBP still increase the response. The addition of more protein cycles would seemingly cause a saturation of the lipid surface, and most of the membrane would be saturated with MBP.

MBP-C1 has a highly positive charge (+19) and depth measurements of the isoforms C1 and C8 have shown that C1 penetrates deeper into a bilayer and binds stronger with both ends of the protein, while the C8 isoform presents greater mobility in the C-terminal end [77]. This would explain why MBP binds so strongly to immobilized vesicles and shed light on the differences seen in the fluorescent patch experiment. MBP-C8 has more flexibility, and residues found in the aqueous region can therefore bind and form new lipid bilayers (Figure 6I-L), whereas MBP-C1 pulls and contracts on the existing membrane (Figure 6AE-H). This in turn can be attributed to its importance at the MDL in holding the inner myelin membrane leaflets tightly together. Furthermore, the areas of double supported membrane patches were found to increase slightly over time after the addition of either P0ct, MBP-C1 or MBP-C8 (Figure 6). For MBP-C1 and MBP-C8, it was observed that the edges of the membrane patches became rougher with extended membrane protrusions. In a previous study, the drug trifluoperazine (TFP), which is known to intercalate into membranes, was observed to induce a remarkable expansion of membrane patches [98]. Previous time-lapse experiments of P0ct indicated no new bilayer formation [73] but formed cavities and lipid accumulation. Conversely, P0ct generated cavities and new bilayers here, suggesting an indistinct mechanism which may direct accumulated lipids into membrane growth. It is important to note that the C-terminal His-tag in the MBP constructs can intervene with protein function and activity [99, 100]. On the other hand, P2 appears to be dependent on P0ct and MBP to bind in larger amounts to lipid bilayers, as increased quantities of P2 rapidly dissociate when it is added alone (Figure 3B). This can be supported by the fluorescent patch experiments, in which a mixture of P0ct and P2 induced new bilayer formation, whereas P2 alone did not generate any new bilayers under the same conditions. These preliminary experiments indicate combined effects from P0ct and P2, and understanding the events taking place in mixtures of these proteins remains a subject for future research.

Regardless of protein-protein interactions, it is evident that P0ct, MBP and P2 bind and stack membranes, producing several diffraction peaks (Figure 4), from which multiple repeat distances were calculated (Supplementary Table 3). The data reveal that most of the protein-lipid aggregates induced by each myelin protein are not purely lamellar or lamellar at all (Supplementary Figure 1). By indexing the peaks, a pure lamellar phase was only discovered in four of the samples tested, while several of the control protein-liposome complexes most likely form a mixture of Im3M/Fd3m at the cost of the lamellar structure. With bicelles, the three proteins induced three bicontinuous phases, Ia3d, Pn3m, and Im3m, respectively, and the micellar cubic phase Fd3m. Cubic phases have been detected several times in biological membranes [101], often in the mitochondria and the endoplasmic reticulum [102, 103]. Furthermore, cubic phases can be induced when cells experience stress, disease, starvation, or viral infections [104, 105], and reactive oxygen species have been found to alter the structural integrity of myelin membranes [106]. Additionally, the lipid PE, which is abundant in biological membranes like myelin [107], is susceptible to forming hexagonal phases [108]. The analysis reflected certain uncertainties for several of the samples and the corresponding Bragg peaks, denoting the need for a more systematic approach in the future. However, as a main result, myelin proteins can induce both lamellar and non-lamellar phases, indicating that P0ct, P2, and MBP can directly influence the physicochemical properties of lipid bilayers. How these direct effects are related to the myelination process, remains a subject for future work.

Given that many of the SAXD samples could not clearly be identified as a cubic phase, and the lattice symmetry might be a mixture of lamellar and cubic phases, we investigated published literature on myelin proteins and Bragg peaks. Several of the calculated repeat distances listed in Supplementary Table 3 can be explained in relation to existing data, illustrated in Figure 7. Knoll *et. al* used neutron scattering to investigate the potential synergistic effects of MBP and P2 in DOPC:DOPS multilamellar vesicles [109], complementing the repeat distances found here when the two proteins are mixed (∼75 Å). The diameter of P2 is roughly 45 Å [27], and the wider d-spacing when P2 is alone harmonizes with a mean membrane thickness of 37.6 Å [73]. In a mixture with either P0ct or MBP, or both, the modelled MDL shrinks by roughly 10 Å (Figure 7). This suggests that P2 must penetrate deeper into the membrane in the presence of P0ct and MBP, or it gets squeezed out of the membrane multilayer by the other proteins. As phase transition experiments revealed that MBP potentially induced the formation of lipid domains, separating protein-bound DMPG and protein-free areas of DMPC, the intermediate states Knoll *et al.* found in lipids containing MBP and P2 [109] likely exist here as well. They found a repeat distance of ∼65 Å that was designated to protein-free lipid domains. Here, a complementary diffraction peak can be found in the bicelle samples at 67 – 70 Å (Figure 7D) that might show the same type of intermediate state of DMPC lipids. X-ray diffraction measurements of mouse sciatic nerves exposed a thickness of 32 - 37 Å for the MDL [110], which is consistent with the width of the intermembrane space found here (Supplementary Table 3). The broader range might be explained by the simplistic model of the MDL examined here, which counts both lipid membrane mixture and proteins, but does not account for all components of the cytoplasmic compartment of native myelin. Another explanation for the Bragg peaks observed, especially with bicelles, can be the induction of different types of lipid crystal structures induced by the myelin proteins. Such structures have been observed before [111, 112] with *e.g.* poly-*L*-lysine, which physicochemically resembles MBP and P0ct.

**Figure 7:**
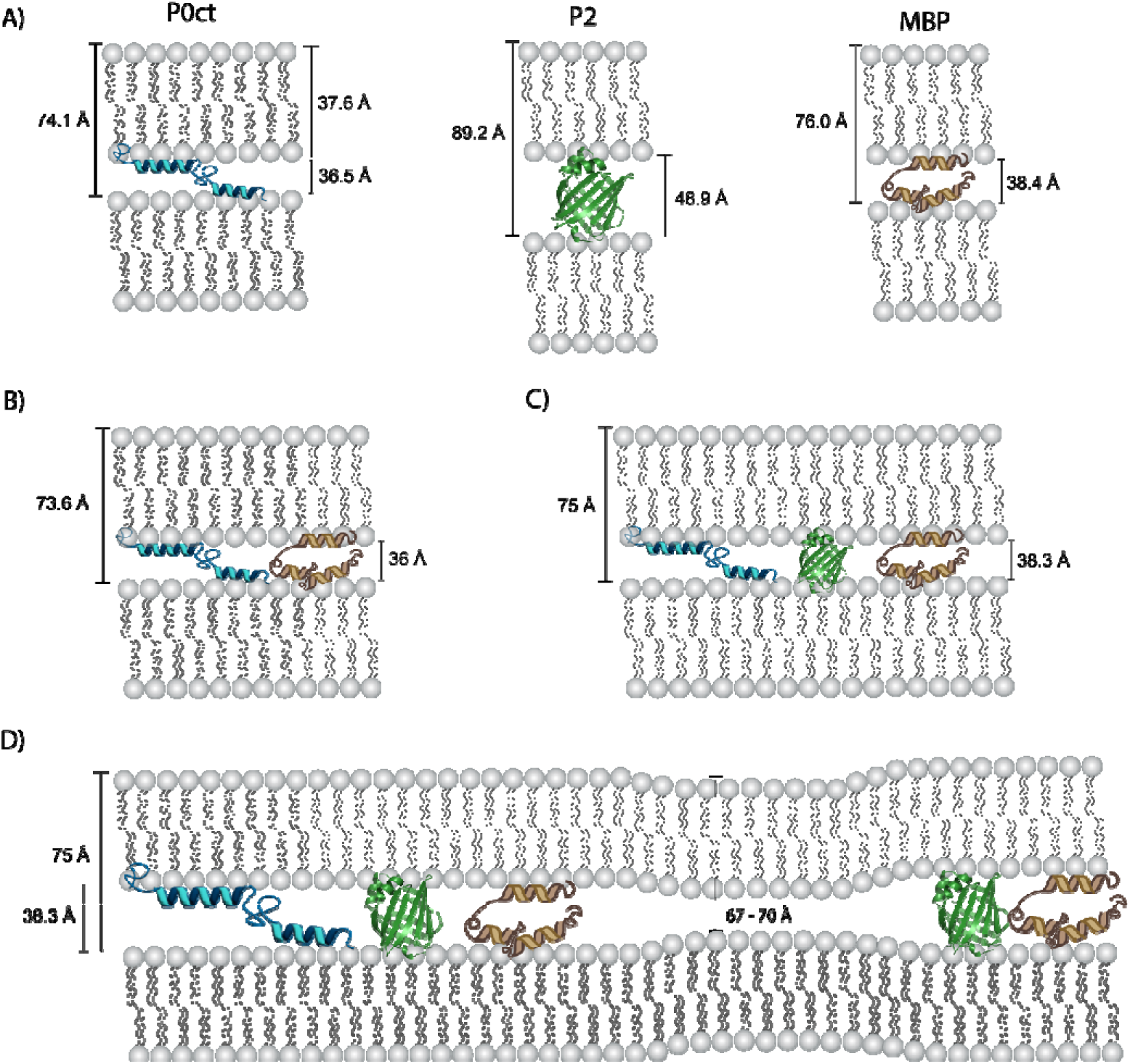
Schematic representation of the repeat distances found in various samples of P0ct (P/L 1:100), P2 (P/L 1:200), and MBP (P/L 1:250) mixed with DMPC:DMPG (1:1) vesicles. The thickness of the SUV membrane has previously been calculated using the MKP method [39] by Hägerbäumer *et al.* [113]. A) Shows lamellar repeat distance when each protein was measured whereas B) and C) list the spacing without and with the addition of P2. D) Potential intermediate state of lipid membrane in protein-bicelle samples. The repeat distances denoted here can be viewed in Supplementary Table 3.

Although SAXD revealed similar peaks and therefore relatable membrane stacking and formation of lipid structures in all mixtures, TEM shows that changes in membrane morphology materialize differently in the different protein-lipid assemblies (Figure 5, Supplementary Figure 7). Measuring the width of two apposing lipid membranes showed a thickness ranging from 12 – 16 nm, which is coherent with the repeat distances found with SAXD, when two membranes are bound together by the proteins investigated, and it is in close range to repeating periods found in sciatic nerve myelin [114]. Myelin consists of layers upon layers of proteolipid membranes in a highly organized fashion, as visualized by electron microscopy [114]. Previous work has shown that P0ct can fuse liposomes and bicelles into larger membrane structures [7, 73]. Although typical bicelle-protein multilayer aggregates were seen when the three proteins were combined, larger stacks of membrane tubes or fibers were present, resembling the ordered fashion of lipid membranes in myelin. These structures appear similar to the MBP-induced clusters of lipid nanotubes observed by cryo-EM [20]. These elongated, thin structures could represent cylindrical or worm-like micelles formed through the reorganization of lipids and detergent, bound together by myelin proteins. Such extended structures can be induced in PC lipids by the addition of small amounts of water or surfactants [115–117], and they were seen previously at high concentrations of P0ct mixed with bicelles [73]. These structures additionally resemble the cylindrical micelles observed earlier for ⍺-synuclein in complex with POPG lipids [118]. Future work will identify the exact interactions and phase behavior of these assemblies. Whether they are relevant for the process of myelination, remains another key question.

## Conclusion

The synergy between MBP and P2 has over the last two decades been explored using atomic force microscopy [54] and neutron diffraction and scattering [109, 119], but investigation of P0ct with MBP and P2 appears to have been lacking until now, despite P0 being the principal PNS myelin protein. Complexes of P0 with PMP22 have been identified on the plasma membrane, while the P0 cytoplasmic tail did not show any interaction with MBP [52]. Regardless, membrane association studies indicated competition between the three proteins during membrane binding. Although P0ct and MBP share similar physical and chemical characteristics, and both bind lipid membranes and condense the intermembrane space between bilayers, they appear to have distinct protein-lipid interplay. P0ct stabilized the membrane, while MBP generated separate lipid domains, producing elevated membrane fluidity. In the CNS, MBP might be part of a molecular boundary between the compact myelin and noncompacted paranodal loops [120], and it is part of different myelin microdomains [88]. MBP might hold a pivotal role in creating discrete lipid rafts within or between compact and noncompact myelin. P2 aids in stacking, while its function appears to be more translucent than P0ct and MBP. However, P2 appears to have a more central role in elongating and fusing lipid membranes, as evidenced by TEM. P2 emerges as a potential contributor in constructing higher-ordered protein-lipid structures. Moreover, SAXD exposed that the three proteins can induce lamellar and cubic phases, which are found in mammalian cells experiencing several stress factors [102, 104]. How this relates to myelin is a question for multidisciplinary experiments in the future. Even though the PNS myelin appears to remain the same without MBP and P2, a reduction in motor nerve velocity, altered lipid composition [121], and reduced myelin thickness have been identified [122]. P2 expression has been associated with thicker myelin sheaths, especially around large motor neuron axons [121, 123]; P2 expression could be a way to increase myelin thickness through synergistically enhancing the combined functions of MBP and P0ct at the MDL. Importantly, missense mutations in both P0ct and P2 have been linked to human CMT neuropathy [15, 33, 124–130], stressing the importance of these myelin MDL proteins for normal myelin structure and function. Using simplistic myelin mimetic models, synergistic effects between P0ct, MBP, and P2 are starting to unravel. Many questions remain on the complex nature of their synergistic effects, but utilizing more complex membrane models – including different lipid compositions and protein mutations – will further aid in our understanding of MDL formation and maintenance and its involvement in human demyelinating disease.

## Acknowledgements

We acknowledge the use of the Core Facility for Biophysics, Structural Biology, and Screening (BiSS) at the University of Bergen, which has received infrastructure funding from the Research Council of Norway (RCN) through NORCRYST (grant number 245828) and NOR–OPENSCREEN (grant number 245922). The help from Dr. Anne Baumann is greatly cherished. In addition, we are grateful to the Molecular Imaging Center (MIC) at the Department of Biomedicine, University of Bergen, for providing access to instrumentation. We acknowledge MAX IV Laboratory for time on the Beamline CoSAXS (Proposal ID 20220497/20200757). Research conducted at MAX IV, a Swedish national user facility, is supported by the Swedish Research Council under contract 2018-07152, the Swedish Governmental Agency for Innovation Systems under contract 2018-04969, and Formas under contract 2019-02496. We gratefully acknowledge the synchrotron radiation facilities and the beamline support at ASTRID2. Parts of the experiments carried out at ASTRID2 were under the support of the EU H2020 MOSBRI research and innovation program. This project has received funding from the European Union’s Horizon 2020 research and innovation programme under grant agreement No 101004806, MOSBRI transnational access proposal ID MOSBRI-2022-137. The assistance of Nykola C. Jones and Søren Vrønning Hoffman at the ASTRID2 synchrotron is highly appreciated. We acknowledge financial support from the University of Bergen (Norway) and several travel grants provided by BioCat, The Norwegian national graduate school in Biocatalysis. The work was supported by a research grant from the Jane and Aatos Erkko Foundation (Finland), and the BIOPROM project is funded by the Research Council of Norway (project number 324877).

## Disclosures

### Author contributions

OCK – conceptualization, data curation, formal analysis, funding acquisition, investigation, methodology, validation, visualization, writing – original draft preparation, writing – review & editing

AR- conceptualization, formal analysis, investigation, methodology, writing – review & editing

MBK - investigation, methodology, resources

AP – investigation, methodology, resources

ØH - investigation

AM – investigation, methodology, resources

SR – investigation, methodology, resources

JSP - conceptualization, methodology, project administration, resources, supervision, validation, writing – review & editing

ACS - conceptualization, methodology, project administration, resources, supervision, validation, writing – review & editing

PK - conceptualization, funding acquisition, investigation, project administration, supervision, validation, writing – original draft preparation, writing – review & editing

### Data availability

The data will be publicly available online at zenodo.org at the time of publication.

## Supplementary Material

Supplementary Figure 1: DSC cooling scans and melting temperatures

Supplementary Figure 2: SRCD spectra of the individual proteins P0ct, MBP, and P2

Supplementary Figure 3: SAXD lattice systems and indexing

Supplementary Figure 4: Kinetic analysis of the SPR data

Supplementary Figure 5: SPR measurements of individual proteins and in sequential order

Supplementary Figure 6: SPR of P0ct, P2 and MBP added in sequence, single graphs

Supplementary Figure 7: TEM control images P0ct, MBP, and P2

Supplementary Figure 8: Preliminary epi-fluorescence patch experiments

Supplementary Figure 9: Turbidity measurements of P0ct, MBP, and P2

Supplementary Table 1: SPR fitting parameters

Supplementary Table 2: Kinetic parameters

Supplementary Table 3: Indexing of SAXD patterns

Supplementary Movie 1: 10 μM P0ct

Supplementary Movie 2: 10 μM MBP-C1-His

Supplementary Movie 3: 10 μM MBP-C8-His

Supplementary Movie 4: 10 μM MBP-C1

Supplementary Movie 5: 2.77 μM MBP-C1 + 2.77 μM P0ct

Supplementary Movie 6: 10 μM P0ct + 10 μM P2

Supplementary Movie 7: 10 μM P0ct + 20 μM P2

Supplementary Movie 8: 13 μM P0ct + 9.87 μM P2

