## Supplementary for "On the synergy between myelin proteins P0, MBP, and P2 in peripheral nerve major dense line formation"

**Supplementary information**

**Supplementary information**

Supplementary Figure 1: DSC cooling scans and melting temperatures

Supplementary Figure 2: SRCD spectra of the individual proteins P0ct, MBP, and P2

Supplementary Figure 3: SAXD lattice systems and indexing

Supplementary Figure 4: Kinetic analysis of the SPR data

Supplementary Figure 5: SPR measurements of individual protein and in sequential order

Supplementary Figure 6: SPR of P0ct, P2 and MBP added in sequence, single graphs

Supplementary Figure 7: TEM control images P0ct, MBP, and P2

Supplementary Figure 8: Preliminary epi-fluorescence patch experiments

Supplementary Figure 9: Turbidity measurements of P0ct, MBP, and P2

Supplementary Table 1: SPR fitting parameters

Supplementary Table 2: Kinetic parameters

Supplementary Table 3: Indexing of SAXD patterns

Supplementary Movie 1: 10 μM P0ct*

Supplementary Movie 2: 10 μM MBP-C1-His*

Supplementary Movie 3: 10 μM MBP-C8-His*

Supplementary Movie 4: 10 μM MBP-C1*

Supplementary Movie 5: 2.77 μM MBP-C1 + 2.77 μM P0ct*

Supplementary Movie 6: 10 μM P0ct + 10 μM P2*

Supplementary Movie 7: 10 μM P0ct + 20 μM P2*

Supplementary Movie 8: 13 μM P0ct + 9.87 μM P2*

* Available at https://doi.org/10.5281/zenodo.12771274

A)

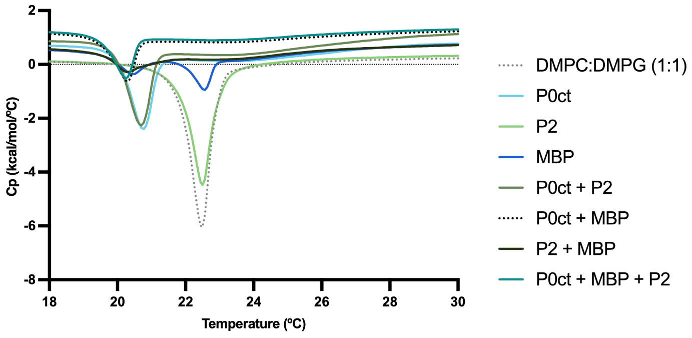

B)

| Mixture | $T_m$ (°C) Cooling | $\Delta T_m$ (°C) Cooling | $\Delta H_{cal}$ Kcal/mol |
|---|---|---|---|
| DMPC:DMPG 1:1 | 22.43 | - | 5.16 |
| P0ct | 20.70 | -1.73 | 1.53 |
| P2 | 22.51 | +0.08 | 4.16 |
| MBP | 20.4, 22.5 | -1.99, +0.07 | 0.66 |
| P0ct + MBP | 20.26 | -2.17 | 0.19 |
| P0ct + P2 | 20.68 | -1.75 | 1.37 |
| P2 + MBP | 20.27 | -2.16 | 0.14 |
| P0ct + P2 + MBP | 20.24 | -2.19 | 0.14 |

**Supplementary Figure 1: DSC cooling scans of A) DMPC:DMPG (1:1) MVLs without and with the addition of native-like protein compositions**. B) The cooling scans' melting temperature ($T_m$), $\Delta T_m$ and $\Delta H_{cal}$.

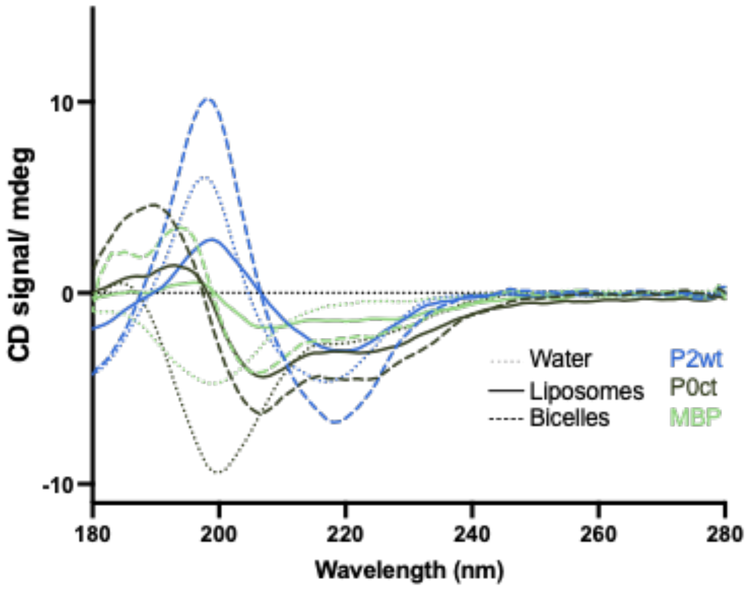

**Supplementary Figure 2: SRCD spectra of the individual proteins P0ct, P2, and MBP in solution (dotted line), in DMPC:DMPG (1:1) SUV liposomes (solid lines), or in bicelles (dashed lines).** The baseline was subtracted from each data set before plotting the spectra. SRCD spectra were measured at +30°C in the wavelength range of 175-280 nm. 5 mM liposomes or bicelles were mixed with each protein right before measuring with a final protein-to-lipid ratio of 1:100 (P0ct), 1:200 (P2), 1:250 (MBP), equivalent to 50 µM P0ct, 25 µM P2, and 20 µM MBP, respectively.

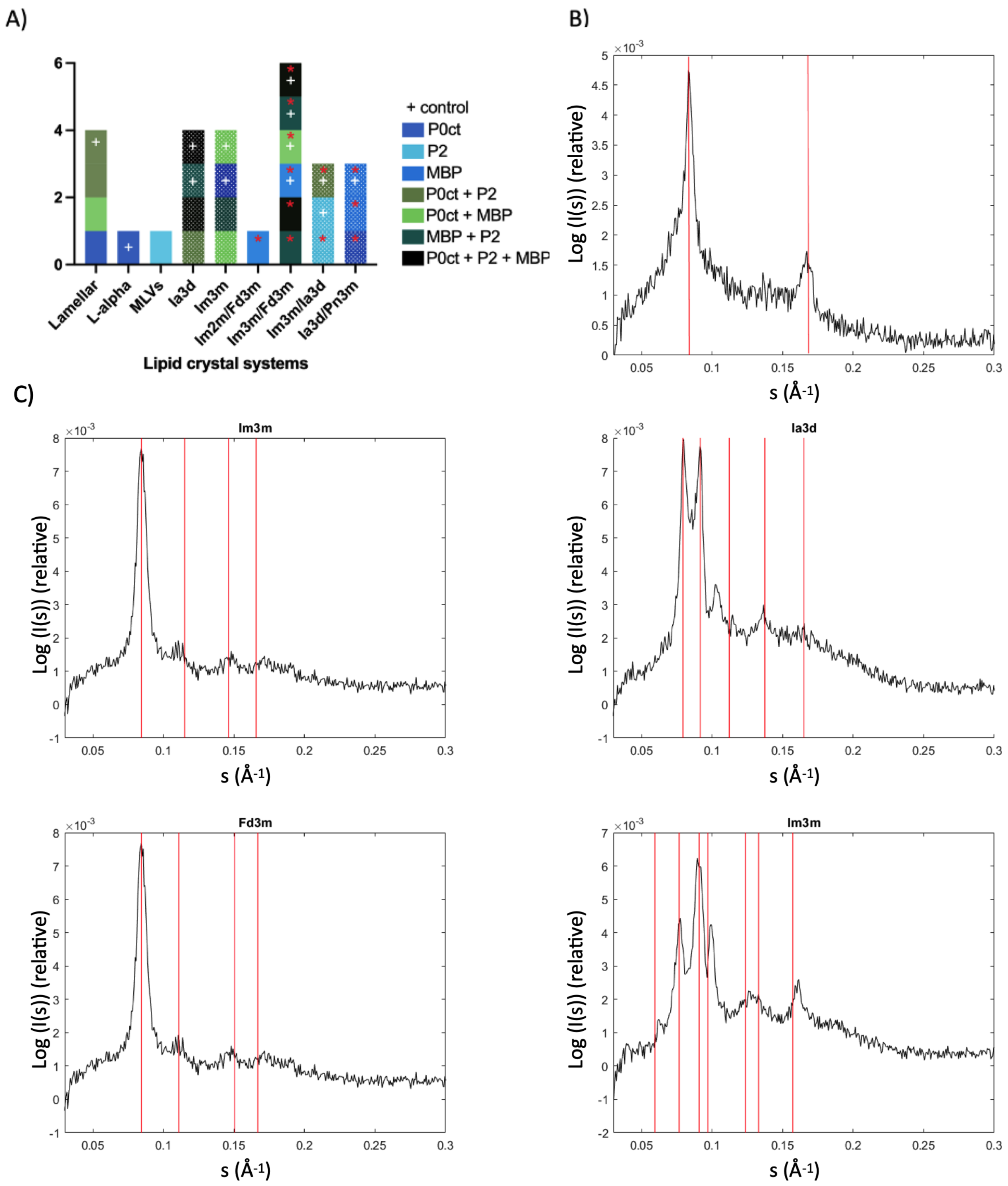

**Supplementary Figure 3: Lattice systems detected in the SAXD measurements of native-like protein compositions and controls**. A) Protein-liposome complexes are shown as filled colored columns, while protein-bicelle mixtures are patterned. Controls are highlighted with a (+), and a red asterisk (*) indicates some uncertainty with the analysis. B-C) Indexing examples of different lattice types. B) shows a lamellar lattice and C) different types of cubic lattice.

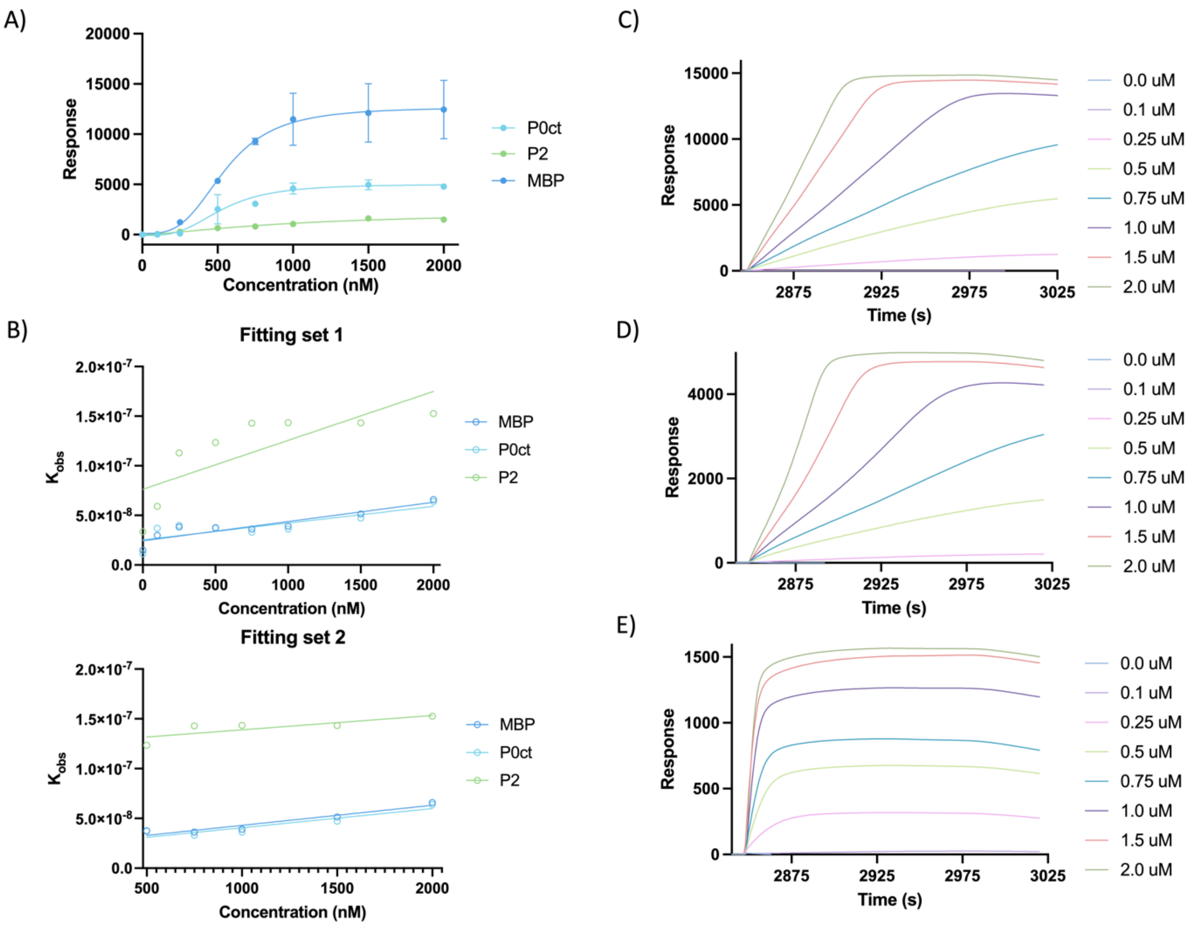

**Supplementary Figure 4: Kinetic analysis of SPR data.** A) Association of each protein with immobilized lipid vesicles using SPR. Error bars represent standard deviation. SPR fitting parameters can be viewed in Supplementary Table 1. Association data of C) MBP, D) P0ct and E) P2 binding to DMPC:DMPG (1:1) LUV vesicles was fitted individually to a one-phase exponential association binding model in GraphPad Prism 10. The derived $k_{obs}$ values were plotted against protein concentration from which $k_{on}$ and $k_{off}$ values were extracted (see Supplementary Table 2) by fitting linear regression functions to the dataset (B – Fitting set 1) and by also omitting all data points before the critical binding concentration, resulting a better linear fit (C – Fitting set 2).

**Supplementary Table 1:** SPR fitting parameters. $A_1$ corresponds to the apparent $K_d$.

| **Sample** | **$R_{hi}$** | **$R_{lo}$** | **$A_1$ (nM)** | **$A_2$** | **$R^2$** |
|---|---|---|---|---|---|
| P0ct | 5029 ± 472.15 | -86.36 ± 298.12 | 526.0 ± 63.35 | 3.171 ± 1.216 | 0.9456 |
| P2 | 2297 ± 768.75 | -19.87 ± 86.40 | 1019 ± 505.43 | 1.421 ± 0.487 | 0.9584 |
| MBP | 12689 ± 989.60 | 117.5 ± 806.11 | 548.5 ± 59.115 | 3.296 ± 1.161 | 0.9400 |

**Supplementary Table 2:** Kinetic parameters derived from each association phase with DMPC:DMPG (1:1) vesicles.

| Sample | Fitting set 1[a] | | | Fitting set 2[a] | | |
|---|---|---|---|---|---|---|
| | $K_{on}$ ($nM^{-1}$ $s^{-1}$)[b] | $K_{off}$ ($s^{-1}$)[c] | $R^2$ | $K_{on}$ ($nM^{-1}$ $s^{-1}$)[b] | $K_{off}$ ($s^{-1}$)[c] | $R^2$ |
| P0ct | $1.683\times10^{-11}$ | $2.539\times10^{-8}$ | 0.6561 | $1.926\times10^{-11}$ | $2.136\times10^{-8}$ | 0.8543 |
| P2 | $4.928\times10^{-11}$ | $7.631\times10^{-8}$ | 0.6214 | $1.441\times10^{-11}$ | $1.245\times10^{-7}$ | 0.6597 |
| MBP | $1.945\times10^{-11}$ | $2.431\times10^{-8}$ | 0.8481 | $2.020\times10^{-11}$ | $2.281\times10^{-8}$ | 0.9216 |

[a] Fitting set 1 contains all data points from the linear fit, whereas all data points below 500 nM were omitted from Fitting set 2.
[b] slope of the linear fit function to $k_{obs(on)}$ vs. [protein]
[c] Y-axis intercept of the linear fit function to $k_{obs(on)}$ vs. [protein]

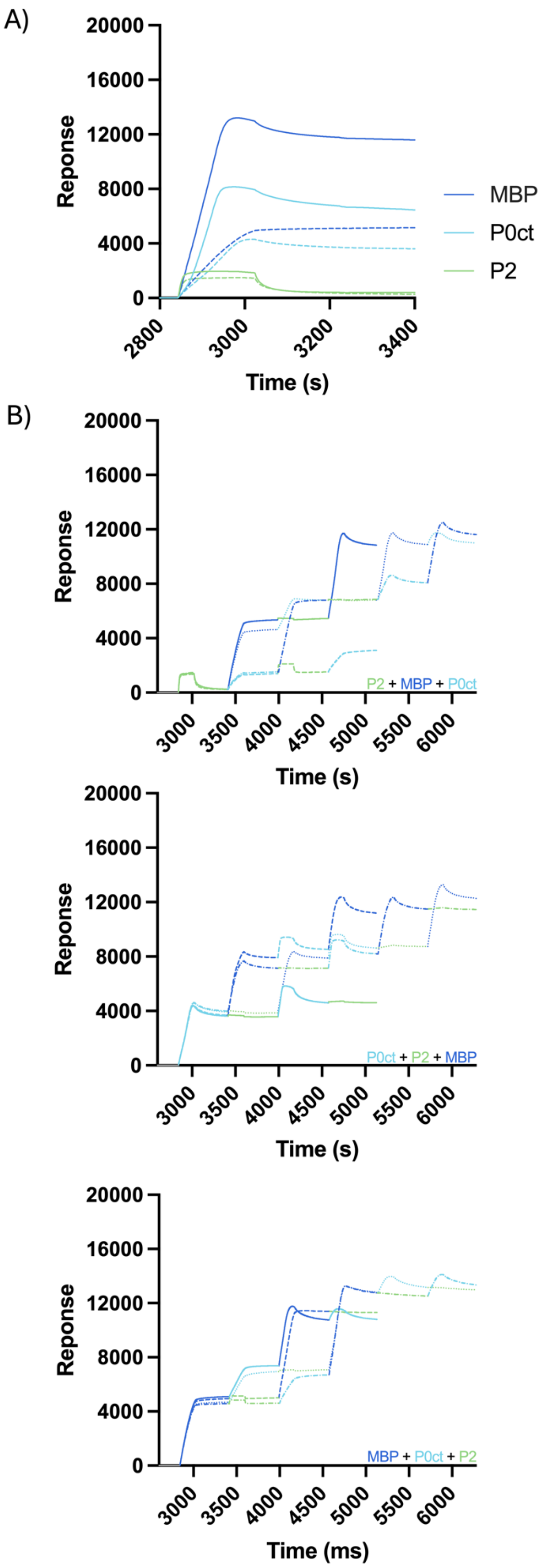

**Supplementary Figure 5: SPR measurements of individual proteins and in sequential order**. A) Dotted lines are from sequence measurements, in which the concentration of MBP and P0ct was reduced to 0.5 µM (dark and light blue dotted line) and P2 concentration was kept at 1.0 µM (green dotted line). Solid lines are all measurements of 1.0 µM protein. B) Proteins in sequence with reduced concentration of MBP and P0ct to 0.5 µM; P2 was kept at 1.0 µM.

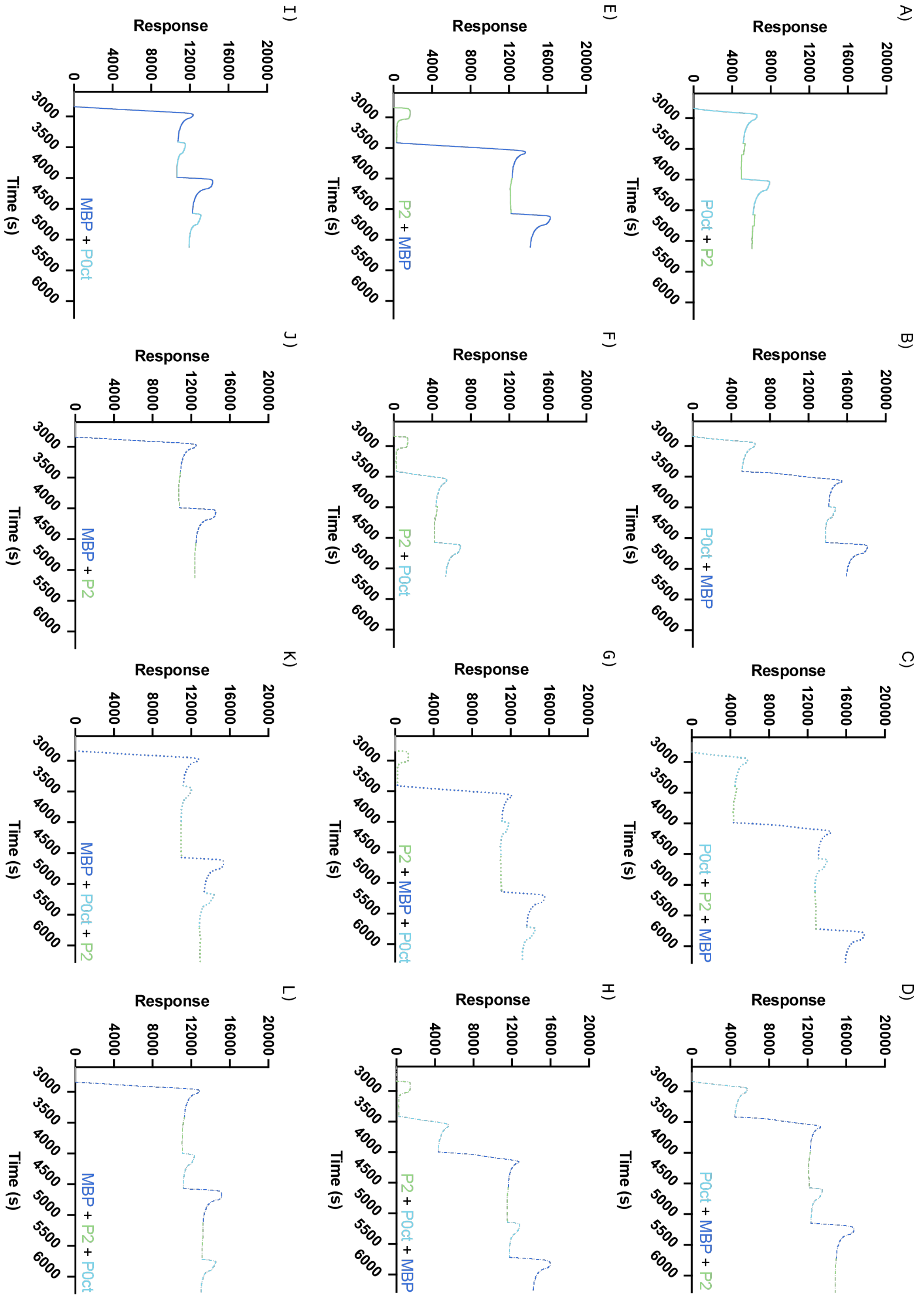

**Supplementary Figure 6: SPR of P0ct, P2 and MBP added in sequence as single graphs**. For easier visualization, SPR measurements shown in Figure 3C-D are shown here as separate graphs. As before: P0ct – light blue, P2 – green, MBP – dark blue. the sequence order is shown in each graph and color-coded. A-D) P0ct as the first protein to be injected, E-H) P2 as the first protein and I-L) MBP as the first protein.

**Supplementary Table 3**: Indexing of the diffraction patterns for each SAXD sample.

| Sample | Phase | lattice parameter (a in Å) |
|---|---|---|
| **3 mM liposomes mixed with:** | | |
| 30 µM P0ct + 15 µM P2 + 12 µM MBP | Im3m/Fd3m is forming at the cost of lamellar phase | $a_{Im3m} = 278.58/a_{Fd3m} = 246.94$ |
| 15 µM P2 + 12 µM MBP | Im3m/Fd3m is forming at the cost of lamellar phase | $a_{Im3m} = 278.58/a_{Fd3m} = 246.94$ |
| 30 µM P0ct + 15 µM P2 | lamellar | $a_L = 75.01$ |
| 30 µM P0ct + 12 µM MBP | lamellar | $a_L = 73.89$ |
| 15 µM P2 | lamellar | $a_L = 86.14$ |
| 50 µM P0ct | lamellar | $a_L = 74.45$ |
| 12 µM MBP | could be Im3m/Fd3m in the molten form | $a_{Im3m} = 282.84/a_{Fd3m} = 250.70$ |
| 16.6 µM P0ct + 16.6 µM P2 + 16.6 µM MBP | Im3m/Fd3m is forming at the cost of lamellar phase | $a_{Im3m} = 278.58/a_{Fd3m} = 246.94$ |
| 25 µM P2 + 25 µM MBP | Im3m/Fd3m is forming at the cost of lamellar phase | $a_{Im3m} = 278.58/a_{Fd3m} = 246.94$ |
| 25 µM P0ct + 25 µM P2 | lamellar | $a_L = 75.01$ |
| 25 µM P0ct + 25 µM MBP | Im3m/Fd3m is forming at the cost of lamellar phase | $a_{Im3m} = 278.58/a_{Fd3m} = 246.94$ |
| 50 µM P2 | unidentified | |
| 50 µM P0ct | lamellar | $a_L = 74.45$ |
| 50 µM MBP | Im3m/Fd3m is forming at the cost of lamellar phase | $a_{Im3m} = 278.58/a_{Fd3m} = 246.94$ |
| **3 mM bicelles mixed with:** | | |
| 30 µM P0ct + 15 µM P2 + 12 µM MBP | Ia3d | $a_{Ia3d} = 192.53$ |
| 15 µM P2 + 12 µM MBP | Im3m | $a_{Im3m} = 262.74$ |
| 30 µM P0ct + 15 µM P2 | Ia3d | $a_{Ia3d} = 192.53$ |
| 30 µM P0ct + 12 µM MBP | Im3m | $a_{Im3m} = 280.68$ |
| 15 µM P2 | mixture of Im3m and Ia3d | $a_{Im3m} = 264.60/a_{Ia3d} = 173.22$ |
| 30 µM P0ct | could be Ia3d/Pn3m | $a_{Ia3d} = 186.60/a_{Pn3m} = 107.73$ |
| 12 µM MBP | could be Ia3d/Pn3m | $a_{Ia3d} = 172.00/a_{Pn3m} = 99.30$ |
| 16.6 µM P0ct + 16.6 µM P2 + 16.6 µM MBP | Ia3d | $a_{Ia3d} = 194.08$ |
| 25 µM P2 + 25 µM MBP | Ia3d | $a_{Ia3d} = 192.53$ |
| 25 µM P0ct + 25 µM MBP | Im3m | $a_{Im3m} = 274.45$ |
| 25 µM P0ct + 25 µM P2 | mixture of Im3m and Ia3d | $a_{Im3m} = 258.52/a_{Ia3d} = 169.24$ |
| 50 µM P2 | mixture of Im3m and Ia3d | $a_{Im3m} = 330.38/a_{Ia3d} = 216.28$ |
| 50 µM P0ct | Im3m | $a_{Im3m} = 303.78$ |
| 50 µM MBP | could be Ia3d/Pn3m | $a_{Ia3d} = 166.08/a_{Pn3m} = 95.89$ |

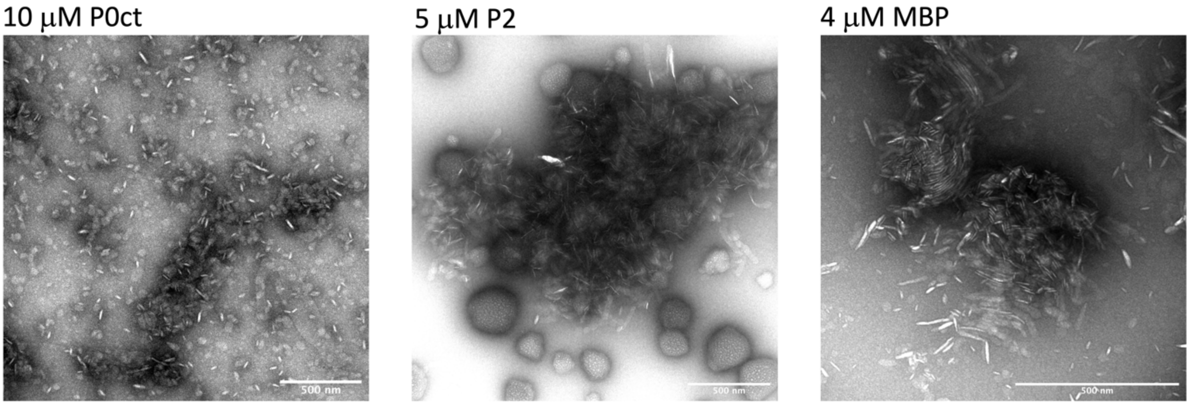

**Supplementary Figure 7: TEM images of the individual proteins:** 10 μM P0ct, 5 μM P2, and 4 μM MBP were mixed with 1 mM bicelles (DMPC:DMPG 1:1) at P/L of 1:100, 1:200, and 1:250, respectively.

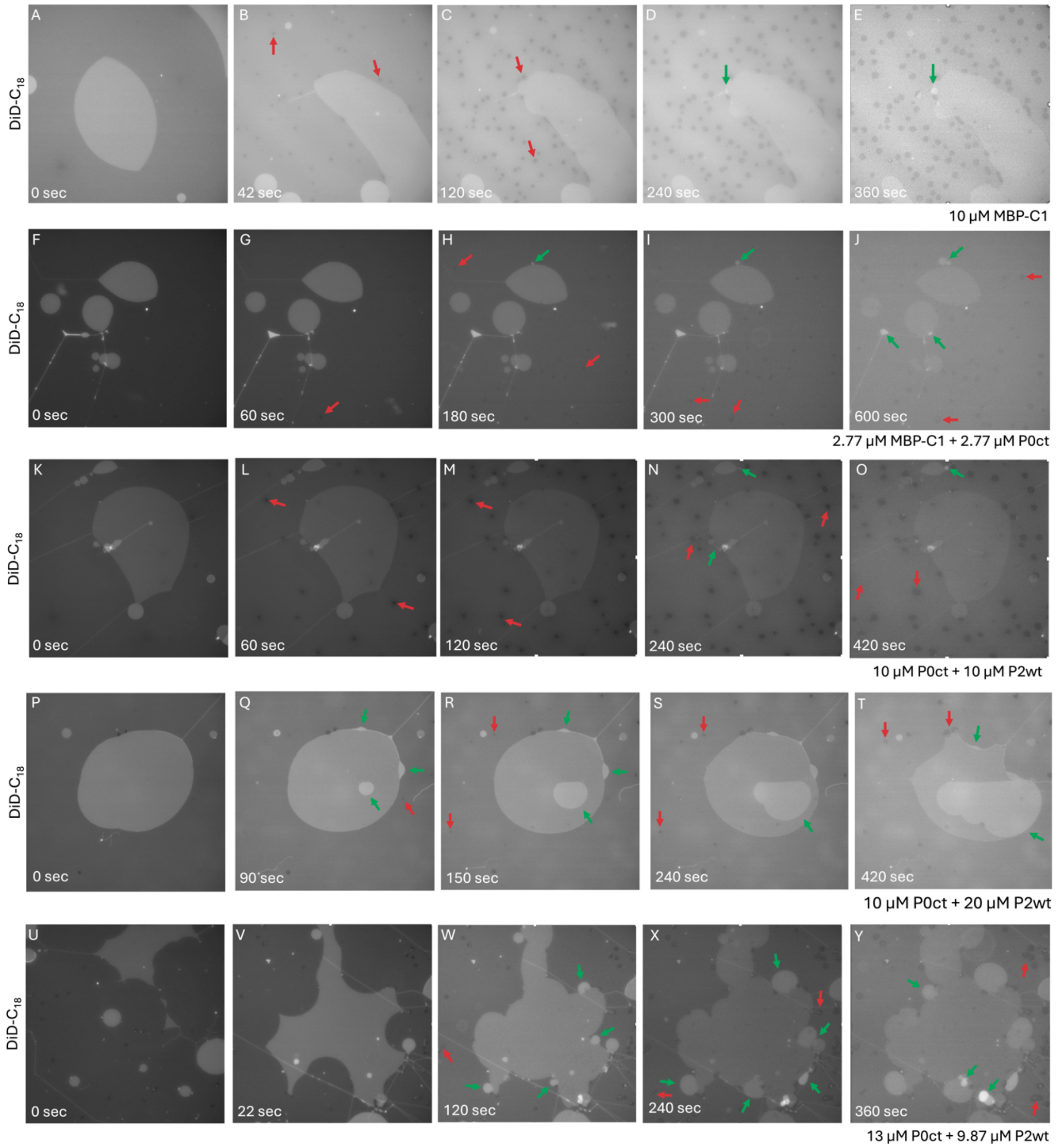

**Supplementary Figure 8: Preliminary fluorescence patch trials of various P0ct, P2 and MBP mixes.** Each experiment was only done once due to the limited amount of protein and needs further examination to be verified. Protein(s) was added to DOPC:DOPS (9:1, 0.05% DiD-C18) and visualized for 15 min. A-E) 10 µM MBP, F-J) 2.77 µM MBP and P0ct, K-O) 10 µM P0ct and P2, P-T) 10 µM P0ct and 20 µM P2, U-Y) 13 µM P0ct and 9.87 µM P0ct. Original movies can be viewed in Supplementary Movie 4-8. With time, the fluorescent bilayers become bleached due to prolonged exposure to light.

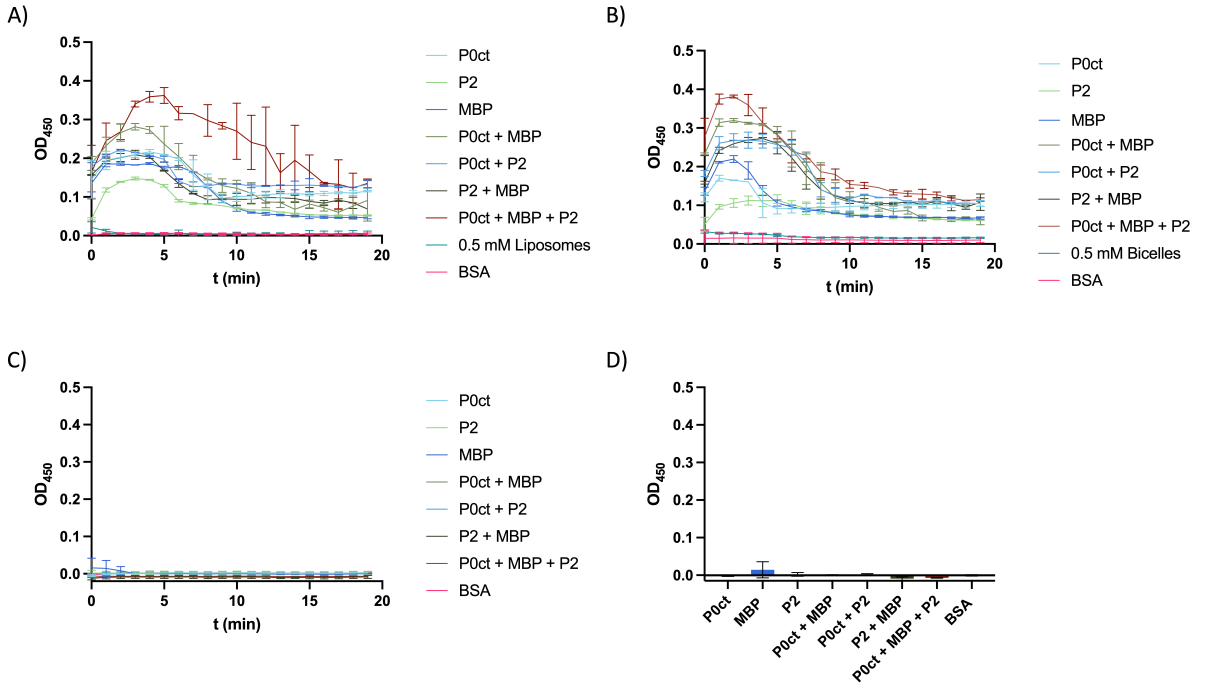

**Supplementary Figure 9: Turbidity measurements of the three proteins P0ct, MBP and P2 in A) 0.5 mM liposomes, B) 0.5 mM bicelles and C) buffer (20 mM HEPES, 150 mM NaCl, pH 7.5) as individual proteins and mixes. D) $OD_{450}$ at t = 1 min for samples measured in HBS.** Protein concentration corresponds to a protein-to-lipid ratio of 1:100 for P0ct, 1:200 for P2, and 1:250 for MBP in all samples. The protein components were added to the aqueous solution containing liposomes, bicelles or only buffer and measured immediately after thorough mixing. Optical density at 450 nm was measured for 20 min at +30 °C. BSA was used as a control with P/L of 1:100. All measurements were done in duplicates and plotted after buffer subtraction.

**Supplementary movies:**

- Supplementary Movie 1: 10 μM P0ct
- Supplementary Movie 2: 10 μM MBP-C1-His
- Supplementary Movie 3: 10 μM MBP-C8-His
- Supplementary Movie 4: 10 μM MBP-C1
- Supplementary Movie 5: 2.77 μM MBP-C1 + 2.77 μM P0ct
- Supplementary Movie 6: 10 μM P0ct + 10 μM P2
- Supplementary Movie 7: 10 μM P0ct + 20 μM P2
- Supplementary Movie 8: 13 μM P0ct + 9.87 μM P2

Available at https://doi.org/10.5281/zenodo.12771274
